# DEAD-box ATPase Dbp2 mediates mRNA release after 3’-end formation

**DOI:** 10.1101/2024.02.17.580811

**Authors:** Ebru Aydin, Jacqueline Böhme, Birte Keil, Silke Schreiner, Bojan Žunar, Timo Glatter, Cornelia Kilchert

## Abstract

mRNA biogenesis in the eukaryotic nucleus is a highly complex process. The numerous RNA processing steps are tightly coordinated to ensure that only fully processed transcripts are released from chromatin for export from the nucleus. Here, we present the hypothesis that fission yeast Dbp2, a ribonucleoprotein complex (RNP) remodelling ATPase of the DEAD-box family, is the key enzyme in an RNP assembly checkpoint at the 3’-end of genes. We show that Dbp2 interacts with the cleavage and polyadenylation complex (CPAC) and localizes to cleavage bodies, which are enriched for 3’-end processing factors and proteins involved in nuclear RNA surveillance. Upon loss of Dbp2, 3’-processed, polyadenylated RNAs accumulate on chromatin and in cleavage bodies, and CPAC components are depleted from the soluble pool. Under these conditions, cells display an increased likelihood to skip polyadenylation sites and a delayed transcription termination, suggesting that levels of free CPAC components are insufficient to maintain normal levels of 3’-end processing. Our data support a model in which Dbp2 is the active component of an mRNP remodelling checkpoint that licenses RNA export and is coupled to CPAC release.

## Introduction

The expression of protein-coding genes depends on the production of functional mRNAs. In eukaryotic cells this involves the formation of the 3’-end of the mRNA by endonucleolytic cleavage and subsequent poly-adenylation (CPA), a prerequisite for the export of mRNAs from the nucleus and their efficient translation in the cytoplasm (Boreikaitė & Passmore, 2023; Gruber & Zavolan, 2019; Rodríguez-Molina & Turtola, 2023). CPA is carried out by the cleavage and polyadenylation complex (CPAC), consisting of cleavage and polyadenylation factor (CPF) and cleavage factors IA and IB (CFIA and CFIB) in yeast, and cleavage and polyadenylation specificity factor (CPSF), cleavage stimulatory factor (CstF) and the mammalian cleavage factors I and II (CFIm and CFIIm) in mammals. Despite differences in complex organisation and the existence of divergent accessory factors, the core components and the general mechanism of CPA are highly conserved (Boreikaitė & Passmore, 2023). First, CPAC is recruited to an elongating RNA polymerase II (RNAPII) (Ahn et al., 2004; Carminati et al., 2023). Components of CPAC then recognize consensus elements contained within the nascent transcript that define the polyadenylation site (PAS), including the canonical AAUAAA signal upstream of the cleavage site and various accessory elements that can be located either up- or downstream (Neve et al., 2017; Porrua & Libri, 2015; Proudfoot, 2011; Rodríguez-Molina et al., 2023; Shi & Manley, 2015). Recognition of the PAS triggers RNA cleavage by the CPAC-associated endonuclease (Ysh1 in yeast, CPSF-73 in humans). The polymerase module then adds a stretch of non-templated adenosines to the newly generated RNA 3’-end to form the poly(A) tail, a process that is controlled by poly(A)-binding proteins that associate with the nascent poly(A) tail (Boreikaitė & Passmore, 2023; Eckmann et al., 2011; Rodríguez-Molina & Turtola, 2023). Importantly, 3’-end processing is tightly coupled to transcription termination: The accessible 5’-PO_4_ end of the downstream cleavage product is a substrate for the 5’-3’ exonuclease Xrn2 (Dhp1 in *S. pombe*), which degrades the nascent RNA until it catches up with RNAPII, thereby helping to displace it from the DNA in a process termed torpedo-mediated transcription termination (Fong et al., 2015; M. Kim et al., 2004; Larochelle et al., 2018; West et al., 2004). The coupling between 3’-end processing and transcription termination is further promoted by the activation of the CPAC-associated phosphatase Dis2/PNUTS-PP1 during PAS recognition. Dis2/PNUTS-PP1 then dephosphorylates the elongation factor Spt5 to slow down transcription in the termination zone, giving the torpedo nuclease an edge over RNAPII (Cortazar et al., 2019; Kecman, Kuś, et al., 2018; Parua et al., 2018).

In addition, there is extensive crosstalk between RNA 3’-end formation and other steps of RNA processing, and the presence of various mRNA biogenesis factors can affect CPAC activity. For instance, splicing has a strong impact on 3’-end processing in mammals: Binding of U1 small nuclear ribonucleoprotein (snRNP) to RNA suppresses cleavage at upstream CPA sites (Kaida et al., 2010; Vagner et al., 2000), whereas interactions of U2 snRNP with CPAC stimulate cleavage activity (Kyburz et al., 2006; Millevoi et al., 2006; Niwa et al., 1990). At the same time, 3’-end processing is modulated by the nuclear RNA surveillance machinery, which can initiate hyperadenylation by the canonical poly(A) polymerase as a signal for RNA decay (Bresson et al., 2015; Chen et al., 2011; Soni et al., 2023; Yamanaka et al., 2010; Zhou et al., 2015).

In contrast to the splicing reaction, in which intronic splice sequences are fully removed, CPA leaves most of the consensus elements that recruit CPAC intact. CPAC components have been found to crosslink to polyadenylated RNA in poly(A) interactome capture experiments in various organisms, including the fission yeast *Schizosaccharomyces pombe* (*S. pombe*) (Baltz et al., 2012; Bao et al., 2018; Beckmann et al., 2015; Castello et al., 2012; Kilchert et al., 2020; Matia-González et al., 2015). A role for poly(A)-binding proteins in preventing re-cleavage of polyadenylated transcripts has been proposed (Viphakone et al., 2008). Evidence from yeast suggests that CPAC components remain with the processed transcripts until mRNA export factors are correctly incorporated into the ribonucleoprotein complex (mRNP) and trigger CPAC release, providing a checkpoint for mRNP assembly that links replacement of CPAC to export competence (Qu et al., 2009). However, the factors that mediate CPAC release upon checkpoint completion have not been identified.

DEAD-box ATPases are good candidates for the implementation of checkpoints that confer directionality to the process of RNA biogenesis. An example for this type of regulation is Dbp5 (DDX19 in humans), which releases export adapters from the freshly exported mRNP upon its activation at the outer face of the nuclear pore complex, thus preventing re-entry of the mRNP into the nucleus (Ledoux & Guthrie, 2011; Lund & Guthrie, 2005; Tran et al., 2007). DEAD-box ATPases belong to an abundant class of proteins that are involved in virtually all aspects of RNA metabolism and are found in all kingdoms of life (Linder, 2006; Putnam & Jankowsky, 2013; Weis & Hondele, 2022). There are at least 33 members of this protein family in humans; of the 19 annotated DEAD-box ATPases in *S. pombe*, most are listed in PomBase as essential for viability (Harris et al., 2022). DEAD-box ATPases are conformationally flexible proteins: In the absence of ATP and RNA, they adopt an open, “off” state, in which their two RecA-like domains, which are connected by a flexible hinge, can have a high degree of structural independence. Upon cooperative binding to ATP and single-stranded RNA, the RecA-like domains reorient to form the closed, “on” state of the active ATPase. ATP hydrolysis leads to RNA release; this can be the rate-limiting step (Hilbert et al., 2009; Ozgur et al., 2015). When bound to a DEAD-box ATPase, the RNA substrate is forced into a kinked conformation that is incompatible with helical structures (Andersen et al., 2006; Del Campo & Lambowitz, 2009; Linder & Lasko, 2006; Sengoku et al., 2006). This mode of RNA binding is central to DEAD-box ATPase function: Distortion of the RNA can result in unwinding of short RNA duplexes (helicase activity) or destabilize RNA-protein interactions, allowing DEAD-box ATPases to remodel mRNPs (RNPase activity) (Donsbach & Klostermeier, 2021; Fairman et al., 2004; Henn et al., 2012; Schwer, 2001; Will & Lührmann, 2001).

Despite a good understanding of the molecular function of DEAD-box ATPases, the high degree of coupling between different RNA processing pathways makes it challenging to link phenotypes that result from DEAD-box ATPase dysfunction – which are often pleiotropic – to a particular RNP remodelling event (Albulescu et al., 2012; Herzel et al., 2018; Moore & Proudfoot, 2009). This also applies to Dbp2 (DDX5 in humans), which has been implicated in many different processes, including transcription, pre-mRNA splicing, nuclear RNA export, RNA decay, RNA-DNA hybrid homeostasis, regulation of liquid-liquid phase-separated compartments, gene looping, RNA silencing, and ribosome biogenesis (Terrone et al., 2022; Weis & Hondele, 2022; Xing et al., 2018). Human DDX5, also known as p68, is involved in transcriptional regulation through its interaction with a histone acetyltransferase, p300/CBP, for which it is also a substrate (Mooney et al., 2010; Rossow & Janknecht, 2003; Wilson et al., 2004). Knockdown of DDX5 in HeLa cells results in the accumulation of unspliced pre-mRNA, and expression levels of DDX5 and its paralog DDX17 influence alternative splicing patterns (Dardenne et al., 2014; Kar et al., 2011; Lin et al., 2005; Terrone et al., 2022). This ability to regulate splicing extends to yeasts, where an (auto-)regulated splicing event in the *DBP2* gene controls Dbp2 expression (Barta & Iggo, 1995; Kilchert et al., 2015). In *S. cerevisiae*, Dbp2 preferentially crosslinks to RNAs that are targets for the Nrd1-Nab3-Sen1 (NNS) termination pathway (Lai et al., 2019; Tedeschi et al., 2018), a budding yeast-specific alternative termination pathway for short stable non-coding RNAs, including small nucleolar RNAs (snoRNAs) (M. Kim et al., 2006; Steinmetz et al., 2001); at NNS target genes, Dbp2 was found to promote transcription termination (Lai et al., 2019). In HeLa cells, read-through transcription was detected at the heat shock gene *HSP70* upon DDX5 depletion, with no observable reduction in PAS cleavage but retention of the *HSP70* transcript at the site of transcription (Katahira et al., 2020). In *Drosophila*, mutants of *Rm62*, the homologue of Dbp2, also fail to release *hsp70* RNA from the site of transcription and display a strong export defect reminiscent of mutations in the core mRNA export receptor *small bristles* (Mex67 in *S. pombe*, NXF1 in humans) (Buszczak & Spradling, 2006). In addition, loss of *DBP2/DDX5* has been found to lead to a redistribution of RNA:DNA hybrids, the total amount of which is increased (Cloutier et al., 2016; Mersaoui et al., 2019; Villarreal et al., 2020).

Here, we report that Dbp2 is present at RNAPII-transcribed genes genome-wide in fission yeast, with a preference for the 3’ end of genes. Using comparative proteomics, we find that Dbp2 interacts with CPAC and RNA export factors. We demonstrate that Dbp2 localizes to cleavage bodies, membrane-less organelles in the nucleus that are rich in CPAC components. Upon loss of Dbp2, 3’-processed, polyadenylated RNAs accumulate in the nucleus and in cleavage bodies, which is accompanied by a depletion of CPAC components from the soluble pool. Under these conditions, cells display an increased tendency to skip PAS sites and a delayed transcription termination, suggesting that the levels of free CPAC components are insufficient to maintain normal levels of 3’-end processing. Our data is consistent with a model in which Dbp2 is a key factor in an mRNP remodelling checkpoint that licenses RNA export and is coupled to CPAC release.

## Results

### Dbp2 is recruited to RNAPII-transcribed genes with a preference for their 3’ ends

To characterize a potential function of Dbp2 in early RNA biogenesis in fission yeast, we carried out chromatin immunoprecipitation of endogenously HTP-tagged Dbp2 followed by sequencing (ChIP-seq). We observed an efficient recruitment of Dbp2 to RNAPII-transcribed genes genome-wide, with a pattern that closely resembled that of an RNAPII-associated RNA processing factor, the SR-like protein Srp2, or of RNAPII itself (Figure 1A). For both Dbp2 and Srp2, the extent of recruitment correlated with amounts of RNAPII at transcribed genes and RNA expression levels, suggesting that Dbp2 is recruited to transcribing RNAPII and/or nascent RNA (Figure 1B and Suppl. Figure 1A and B).

**Figure 1:**
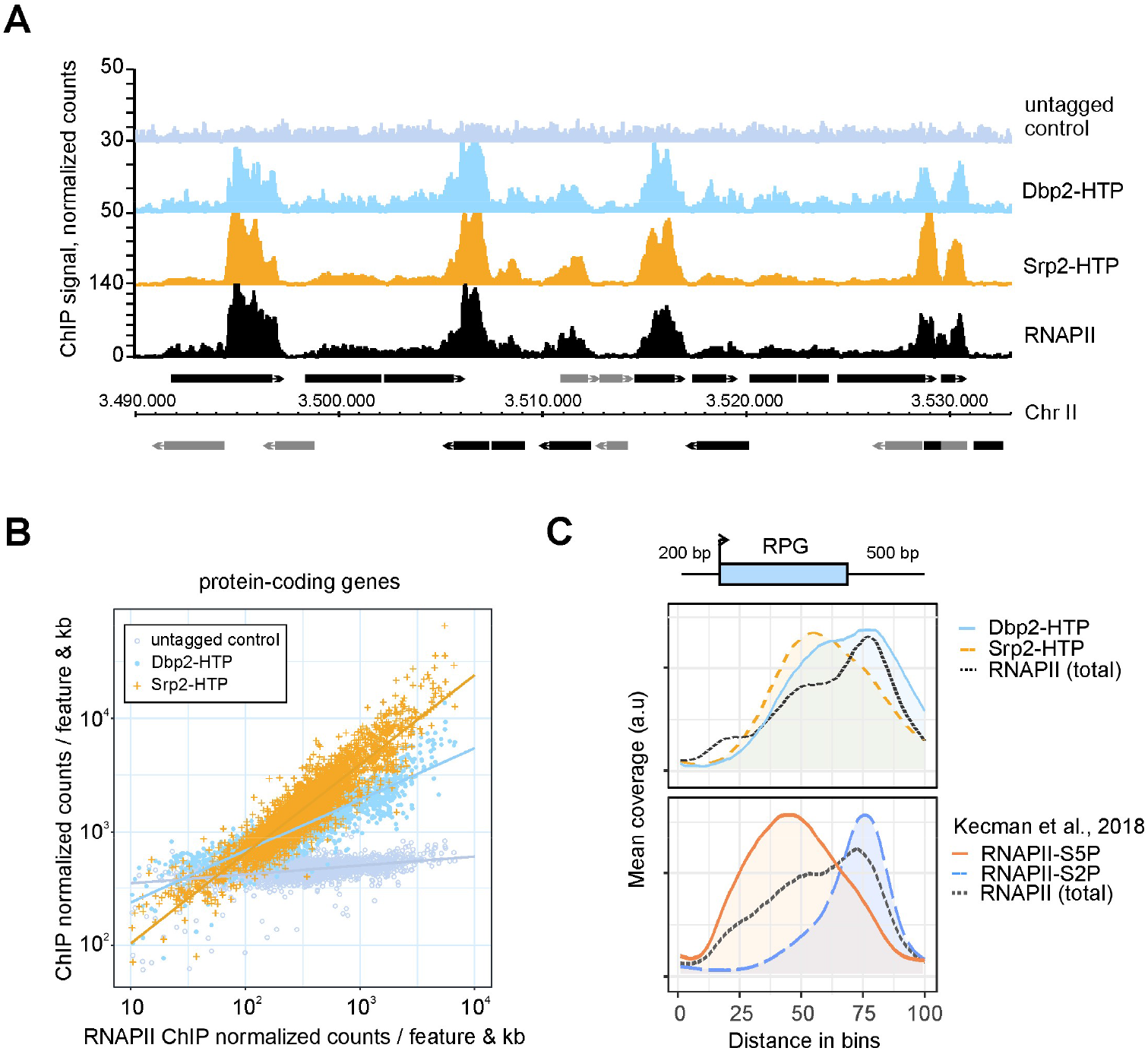
Dbp2 is recruited to RNAPII-transcribed genes genome-wide. **A** Representative ChIP-seq traces of Dbp2-HTP and Srp2-HTP association with chromatin across a region of *S. pombe* chromosome II. An untagged wild type was included as control. RNAPII-ChIP signal for an isogenic wild type is shown for reference (α-Rpb1, 8WG16) (Kecman et al., 2018; GEO: GSE111326). Positions of protein-coding genes and non-coding genes are indicated in black and grey, respectively. **B** Integrated counts of Dbp2-HTP and Srp2-HTP ChIP-seq signal across protein-coding genes relative to RNAPII, given as average counts per feature and kb * 1,000,000 (n = 3). Only genes with an RNAPII occupancy > 10 normalized counts per feature and kb per million were included. Trendlines were fitted using linear regression. RNAPII ChIP data from Kecman et al., 2018; GEO: GSE111326 (n = 2). **C** Metagene analysis of mean ChIP-seq coverage of Dbp2-HTP and Srp2-HTP (n = 3), and total RNAPII (α-rpb1 (8WG16) in Dbp2-3myc; n = 2) (upper panel) across ribosomal protein genes (RPGs) including 200 bp upstream and 500 bp downstream of the annotated transcription units. Mean coverage given in arbitrary units; to compensate for differences in ChIP capability, mean coverage was adjusted by a constant scaling factor for an easier comparison of curve shapes. Non-scaled curves with confidence intervals are provided in Suppl. Figure 1C and D. Total Rpb1, Rpb1-S5P, and Rpb1-S2P ChIP for an isogenic wild type (n = 2) are included as reference (lower panel; data from Kecman et al., 2018; GEO: GSE111326). Schematic of the gene above the plot corresponds to an RPG of median length. Within the metagene, the positions of transcription start and end sites are distributed around the given coordinate because of the varying feature compression depending on gene length.

To determine the recruitment pattern across transcription units, we generated metagene plots comparing the distribution of Dbp2-HTP, Srp2-HTP, and RNAPII along ribosomal protein genes (RPGs) (Figure 1C). We focused on RPGs because they are highly expressed and are characterized by a high ChIP-seq signal above background. To be able to differentiate between stages of transcription (initiating, elongating, and terminating RNAPII), we compared to publicly available ChIP-seq data profiling different modifications of the C-terminal domain (CTD) of Rpb1, the largest subunit of RNAPII (Kecman, Heo, et al., 2018) (Figure 1C, lower panel); these modifications are added and removed as RNAPII transcription progresses and mark specific stages of the transcription cycle, with serine 5 phosphorylation (S5P) and serine 2 phosphorylation (S2P) characteristic for initiating and terminating RNAPII, respectively (Eick & Geyer, 2013; Sanso & Fisher, 2013). The mean ChIP-seq coverage of both Dbp2 and Srp2 increases gradually along gene bodies. However, while the Srp2-HTP coverage peaks within the gene body and then decreases towards the end, the coverage for Dbp2-HTP continues to rise and reaches its highest point after the CPA site, coinciding with the RNAPII-S2P peak (Figure 1C, compare upper and lower panels). Our data are compatible with a function for Dbp2 as a basal regulator of RNAPII transcription or co-transcriptional RNA processing, particularly at the 3’ end of genes.

### Comparative interaction profiling of Dbp2 and Srp2 reveals their association with RNAPII at different stages of the transcription cycle

We next sought to identify direct protein-protein interactors of Dbp2. Initial attempts with stringent purification protocols did not result in the co-purification of significant amounts of proteins or protein complexes with Dbp2-HTP. This behaviour is reminiscent of other DEAD-box ATPases such as Dbp5, whose interactions with its regulators have been described as weak and transient (Montpetit et al., 2011). To stabilize transient interactions, we resorted to mild protein-protein crosslinking and treated cells with formaldehyde prior to lysate preparation (0.01% formaldehyde for 10 min). Based on the ChIP-seq experiments, we anticipated co-purification of the transcription machinery along with the major co-transcriptional RNA processing complexes, including capping factors, the spliceosome, and the 3’ end processing machinery. To be able to assess whether co-purification of protein complexes with Dbp2-HTP reflected a preferential association with Dbp2 over other proteins contained within the same compartment, we also purified the *bona fide* RNAPII interactor Srp2-HTP as a reference using the same conditions (Figure 2A). Co-purifying proteins were quantified by shotgun proteomics (Supplemental Table S1). In agreement with the ChIP-seq results, Dbp2-HTP and Srp2-HTP co-purified components of the RNAPII holoenzyme complex; amounts of co-purifying RNAPII were similar in both cases (Figure 2B and Suppl. Fig. 2A). A notable exception was Rpb8, which co-purified more readily with Srp2-HTP, potentially indicating a direct protein-protein interaction (Figure 2B and Suppl. Fig 2A); results of interaction modelling with AlphaFold2 multimer are consistent with this possibility (Suppl. Fig 2B). Known interactors of Srp2, including Srp1 and the SR protein kinase Dsk1 (Tang et al., 2002), were significantly enriched in the Srp2-HTP purification (Figure 2B and Suppl. Fig. 2A). The most highly enriched protein in the purification of Dbp2-HTP was the type I protein arginine N-methyltransferase (PRMT1) Rmt1 (Figure 2B and Suppl. Fig. 2A). PRMT1 proteins have a strong preference for methylating RGG/RG motifs, multiple copies of which are present in the C-terminus of Dbp2, and the human Dbp2 orthologue DDX5 is a known target of arginine methylation (Mersaoui et al., 2019; Thandapani et al., 2013).

**Figure 2:**
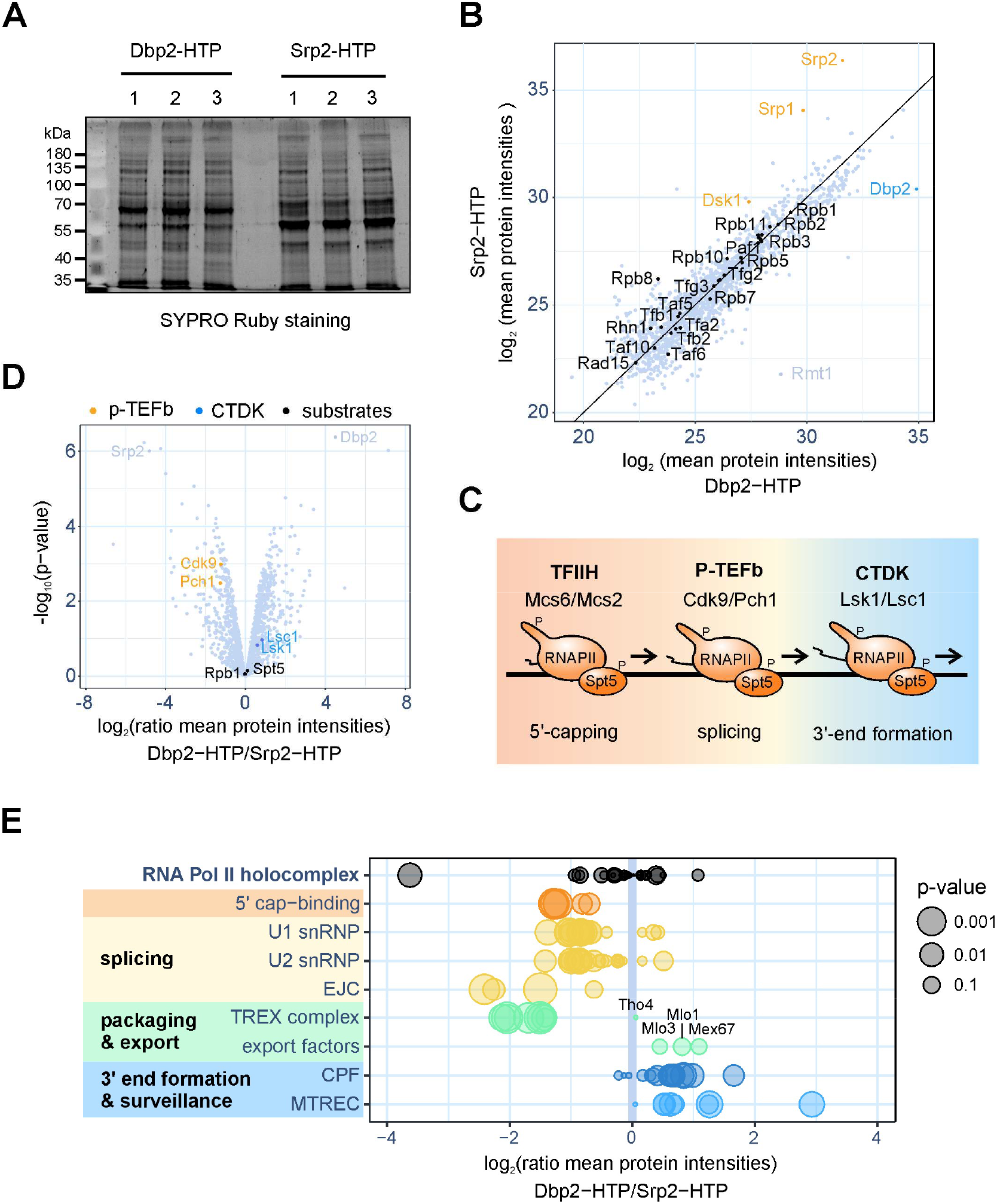
Comparative protein interaction profiling of Dbp2 and Srp2. **A** Eluates of cross-linking HTP purifications of Dbp2-HTP and Srp2-HTP were resolved on SDS-PAGE and stained with SYPRO Ruby. The experiment was carried out with three biological replicates. **B** Mass spectrometry (MS) analysis of the comparative interaction profiling of Dbp2 and Srp2. Mean protein intensities (log_2_) recovered in the Dbp2-HTP and Srp2-HTP purifications (n = 3). Components of the RNAPII holocomplex (GO:0016591) are marked in black, Srp2 and known interactors in orange, and Dbp2 in light blue. **C** Schematic of the CDK-dependent transcription cycle in fission yeast. In the presence of its cognate cyclin, Mcs2, the TFIIH-associated CDK Mcs6 phosphorylates S5P of the RNAPII CTD and Spt5 at the promoter to recruit capping factors. The activity of P-TEFb and its associated CDK/cyclin pair Cdk9/Pch1 is linked to promoter release and splicing. A second S2P-specific CDK complex, CTDK, promotes cleavage and polyadenylation and transcription termination at the 3’ end of genes. Adapted from Sanso & Fisher, 2013. **D** Relative enrichment of cyclin-dependent kinase (CDK) complexes co-purifying with Dbp2 and Srp2. In the volcano plot, p-values (−log10, moderated Student’s t-test) are plotted against the relative enrichment of proteins in the purification of Dbp2-HTP relative to Srp2-HTP based on mean protein intensities (log2) (n = 3). The P-TEFb- and CTDK-associated CDK/cyclin pairs are marked in yellow and blue, respectively. The CDK substrates Rpb1 and Spt5 are marked in black. E Relative enrichment of various RNA processing complexes co-purifying with Dbp2 and Srp2 based on mean protein intensities (log_2_) (n = 3). The size of the circles reflects P-values (moderated Student’s t-test). snRNP – small nuclear ribonucleoprotein. EJC – exon junction complex. CPF – mRNA cleavage factor complex. MTREC – Mtl1-Red1 core. Annotations were retrieved from Ensemble Fungi using the following GO terms: RNAPII holocomplex - GO:0016591; U1 snRNP - GO:0005685; U2 - GO: snRNP – GO:0005686; mRNA cleavage factor complex - GO:0005849. Export factors: Mex67, Mlo1 and Mlo3. Volcano plots where the individual proteins are labelled are provided in Suppl. Figure 2C and D.

Next, we analysed the relative enrichment of RNA biogenesis factors. The modest 5’ to 3’ shift in the chromatin association patterns of Srp2-HTP and Dbp2-HTP had already suggested a temporal offset in their recruitment to transcribing RNAPII (Figure 1C). This assumption is further supported by the comparative interaction profiling data: The cyclin-dependent kinases (CDKs) that are responsible for the placement of the phospho-marks on RNAPII and Spt5 that direct RNA processing are known to be recruited in a sequential manner (Sanso & Fisher, 2013) (Figure 2C). P-TEFb (Cdk9/Pch1 in *S. pombe*), an early RNAPII-S2P kinase with strong links to pre-mRNA splicing that has been implicated in the transition from initiation to productive transcription elongation, was significantly enriched in the Srp2-HTP purification. In contrast, the late S2P kinase complex CTDK (Lsk1/Lsc1) preferentially co-purified with Dbp2-HTP (Figure 2D). Only very low levels of TFIIH were detected in both purifications, in comparable amounts, with the associated S5P kinase complex Mcs6/Mcs2 not co-purifying at all. We conclude that Srp2 and Dbp2 are recruited to transcribing RNAPII after promoter clearance. This is consistent with the ChIP-seq profiles, where chromatin-associated Srp2 and Dbp2 show little overlap with the promoter-associated RNAPII peak (Figure 1C).

### Dbp2 preferentially associates with 3’-end formation factors and the nuclear RNA surveillance machinery

In agreement with the proposed function of Srp2 as a splicing regulator, splicing factors were enriched in the Srp2-HTP purification, as were cap-associated proteins and the exon junction complex (EJC; Figure 2E and Suppl. Figure 2C and D). The TREX complex, an RNA packaging complex that couples mRNA transcription, processing and nuclear export (Jimeno et al., 2002; Sträßer et al., 2002), was significantly enriched in the Srp2-HTP purification, with the exception of Tho4, the RNA export adapter homologous to the human TREX component ALYREF/THOC4 (Suppl. Figure 2D). Notably, *S. pombe* possesses a second ALYREF/THOC4 protein, Mlo3, which was modestly enriched in the Dbp2-HTP purification, along with Mlo1 (homologue of human SARNP/CIP29) and the mRNA export receptor Mex67 (Figure 2E and Suppl. Figure 2D). 3’-end processing factors preferentially co-purified with Dbp2-HTP, in agreement with its presence during the late steps of the transcription cycle. For one representative CPAC component, Msi2, the interaction was validated by non-crosslinking co-immuno-precipitation (Suppl. Figure 2F). In addition, components of the exosome targeting factor MTREC were more enriched in the Dbp2-HTP purification (Figure 2E and Suppl. Figure 2D).

### R-loop-associated proteins are not overrepresented in the Dbp2-HTP purification

Because orthologues of Dbp2 have been implicated in the resolution of RNA:DNA hybrids in the context of transcription-associated R-loops (Mersaoui et al., 2019; Polenkowski et al., 2023; Tedeschi et al., 2018; Yu et al., 2020), we also screened for known R-loop interactors. RNase H enzymes (Rnh1 and Rnh201/202/202) and DNA topoisomerases (Top1, Top2, Top3, Rec12), proteins typically associated with R-loops, were absent from both purifications. R-loop interactomes are also rich in RNA helicases and single-stranded DNA-binding proteins, which coat the displaced DNA strand (Cristini et al., 2018). Proteins of both classes co-purify with Srp2-HTP and Dbp2-HTP, with no obvious preference for either protein; the RNA/DNA helicase Sen1, for example, a known hybrid-dissolving enzyme (H. D. Kim et al., 1999; Mischo et al., 2011), was enriched 1.03-fold in the Dbp2-HTP purification. The histone chaperone FAcilitates Chromatin Transcription (FACT) complex – another known R-loop regulator (Cristini et al., 2018; Herrera-Moyano et al., 2014) – was modestly enriched in the Dbp2-HTP purification (Suppl. Figure 2E). In comparison, RSC, a SWI/SNF family chromatin remodelling complex with essential roles in transcription regulation and the maintenance of nucleosome-depleted regions (Monahan et al., 2008; Yague-Sanz et al., 2017), had a strong preference for co-purifying with Dbp2-HTP, with the RSC-specific essential subunits Snf21, Rsc7 and Rsc9 particularly significantly enriched (Suppl. Figure 2E). The histone lysine acetyltransferase Rtt109, on the other hand, which has structural similarities to human p300, a known interactor of human DDX5 (Bazan, 2008; L. Wang et al., 2008), was not detected in any of the purifications.

### Dbp2 localizes to cleavage bodies

MTREC/PAXT components were among the most highly enriched proteins in the Dbp2-HTP purification. MTREC is not a general component of the transcription machinery but is specifically recruited to exosome target genes by the YTH-domain protein Mmi1, which recognizes TNAAAC motifs on nascent RNA and primes transcripts for decay by the nuclear exosome (Harigaya et al., 2006; Kilchert et al., 2015; Tashiro et al., 2013; Yamashita et al., 2012). We observed no preferential recruitment of Dbp2 to Mmi1-regulated genes (Suppl. Figure 3A). However, MTREC components and 3’-end formation factors co-localize to discrete nuclear bodies in mitotically dividing fission yeast (Egan et al., 2014; Shichino et al., 2020; Sugiyama et al., 2012, 2013; Sugiyama & Sugioka-Sugiyama, 2011; Yamanaka et al., 2010). There is a compositional overlap between these “nuclear exosome foci” and the cleavage bodies described in human cells, and it has been suggested that these structures are analogous (L. Li et al., 2006; Schul et al., 1996; Sugiyama & Sugioka-Sugiyama, 2011). For the sake of brevity, we will refer to MTREC-containing foci as “cleavage bodies” throughout this work. Having found Dbp2 purifications to be enriched for many proteins associated with cleavage bodies in *S. pombe*, we wondered whether Dbp2 would also be present in these granules. A genomic C-terminal fusion of Dbp2 to GFP has been reported to localize to the nucleolus (Matsuyama et al., 2006), which we also observe (Figure 3A and C). In addition, Dbp2-GFP localizes to punctate structures in the nucleus, which are often adjacent to the nucleolus or the nuclear rim. To determine whether these foci correspond to cleavage bodies, we crossed Dbp2-GFP to strains carrying C-terminal tdTomato tags on the MTREC components Red1 or Pab2. Dbp2-GFP signal within foci almost universally overlapped with the Red1-tdTomato signal, which exclusively localized to foci (Figure 3A and B). In contrast, Pab2-tdTomato – while also being concentrated in foci – is present throughout the nucleoplasm and concentrated at the nucleolar rim, but mostly excluded from the nucleolus itself. Again, the punctate Pab2-tdTomato signal colocalized with Dbp2-GFP foci (Figure 3C and D), indicating that Dbp2 is a component of cleavage bodies. We note that while Dbp2 and Pab2 predominantly localize to different, non-overlapping compartments (nucleolus and nucleoplasm, respectively), they intersect in cleavage bodies, which preferentially occur at the interface between both compartments (Figure 3C and D).

**Figure 3:**
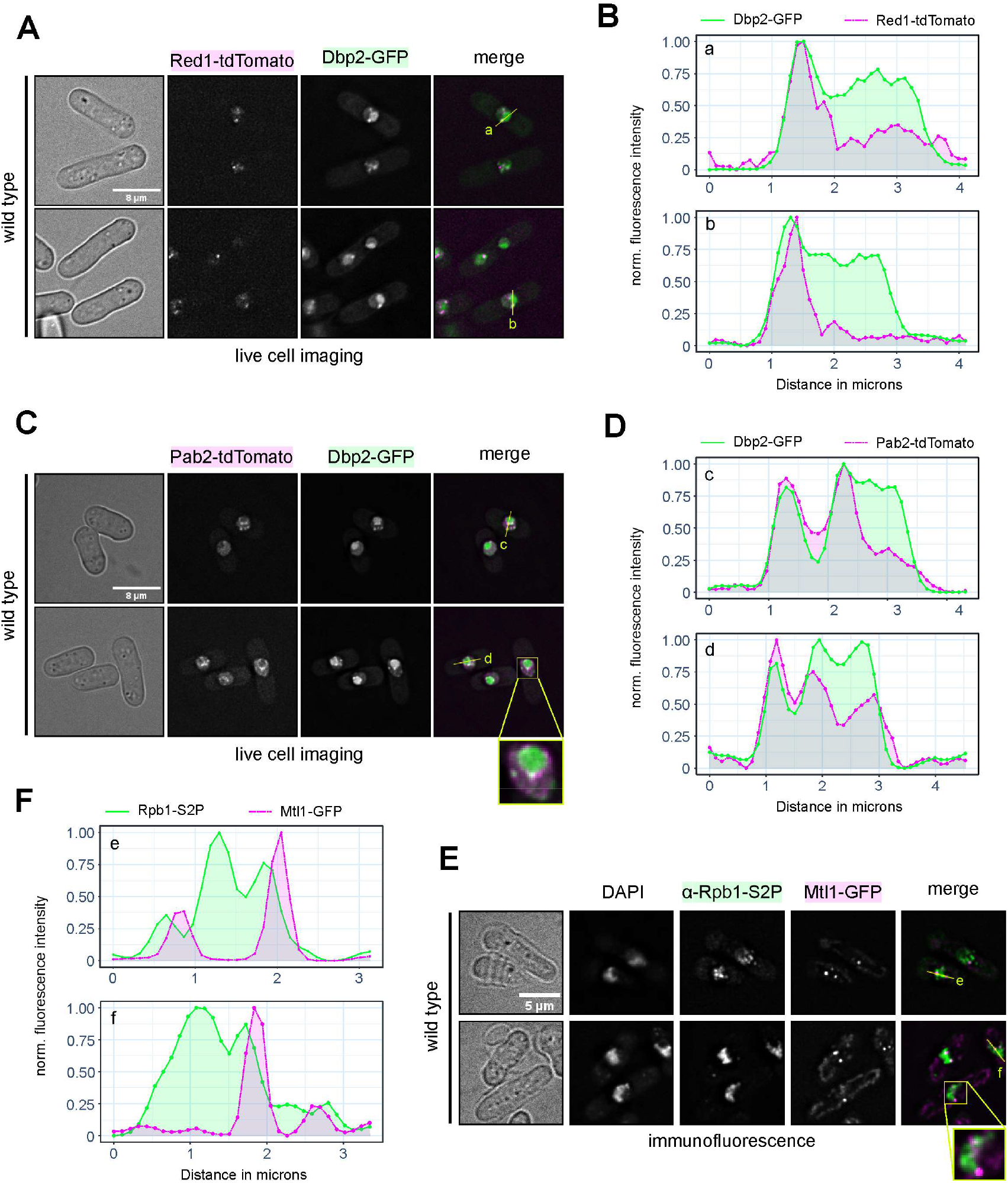
Dbp2 is a component of cleavage bodies. **A** Live cell imaging of genomically tagged Dbp2-GFP and the MTREC component Red1-tdTomato as a marker for cleavage bodies. Cells were grown in YES at 30°C, pelleted and resuspended in EMMG for imaging on poly-lysine-coated coverslips. The merged channel shows Dbp2-GFP in green and Red1-tdTomato in magenta. Fluorescence intensity profiles were generated along the yellow lines and are shown in B. Images are representative of three independent experiments. **B** Fluorescence intensity profiles across cleavage bodies and the adjacent nuclear area as indicated in A. Green line corresponds to the Dbp2-GFP signal, dashed magenta line to the Red1-tdTomato signal. Fluorescence intensities were measured with Fiji and normalised to a 0-1 range. **C** and **D** Live cell imaging and fluorescence intensity profiles of genomically tagged Dbp2-GFP and the nuclear poly(A)-binding protein Pab2-tdTomato as a marker for cleavage bodies as in A and B. Images are representative of three independent experiments. **E** Immunofluorescence against Rbp1-S2P in a strain where MTREC component Mtl1 was genomically tagged with GFP. Cells were grown in YES at 30°C and fixed with formaldehyde directly in the growth medium. The merged channel shows Rpb1-S2P in green and Mtl1-GFP in magenta. Fluorescence intensity profiles were generated along the yellow lines and are shown in F. Images are representative of two independent experiments. Enlarged versions of the merged image without the yellow markings are provided in Suppl. Fig. 3B. **F** Fluorescence intensity profiles across cleavage bodies and sites of active transcription as indicated in E. Green line corresponds to the Rpb1-S2P signal, dashed magenta line to the residual Mtl1-GFP signal. Fluorescence intensities were normalised to a 0-1 range. Images are representative of two independent experiments.

### Cleavage bodies are unlikely to represent sites of co-transcriptional RNA cleavage

Because cleavage bodies harbour factors involved in co-transcriptional RNA processing, we wondered whether they would be associated with highly active sites of transcription. We carried out immune-fluorescence microscopy with an antibody against S2P-modified Rpb1 to detect transcriptionally active RNAPII in a strain where the MTREC component Mtl1 was C-terminally tagged with GFP. Rpb1-S2P yielded a granular staining that overlapped with the DNA-rich crescent-shaped nucleoplasmic compartment counterstained with DAPI (Figure 3E). While cleavage bodies were frequently detected adjacent to an area of higher Rpb1-S2P density, this was largely limited to the surface of the RNAPII transcriptionally active compartment, particularly at the inner arc of the crescent shape, which corresponds to the interface with the nucleolus (Figure 3E and F, and Suppl. Figure 3B). Overall, the majority of RNAPII transcription hot spots were not associated with a cleavage body, and Mtl1-GFP foci did not usually coincide with a peak in Rpb1-S2P signal. We deduce that cleavage bodies, while containing components of the 3’-end formation machinery, are unlikely to represent the sites where the bulk of RNA cleavage and polyadenylation occurs, which is coupled to RNAPII transcription and takes place on the nascent transcript (Larochelle et al., 2018; Rodríguez-Molina et al., 2023). While we cannot exclude that CPAC components within cleavage bodies can still be active on RNAs that have been released from chromatin before processing had completed, we conclude that cleavage bodies most likely represent storage compartments for 3’-end processing factors.

### In the absence of Dbp2, poly(A)+ RNA is retained in the nucleus

Our data indicated that Dbp2 interacts with the 3’-end formation machinery and RNA export factors, suggesting that it might bridge these two processes. We hypothesized that Dbp2 – as a DEAD-box ATPase and potential RNPase – could be involved in CPAC release from processed transcripts as part of the mRNP assembly checkpoint that has been proposed to assess export competence (Qu et al., 2009). To evaluate whether Dbp2 was involved in this process, we generated an inducible depletion strain. Because Dbp2 was resistant to depletion via an auxin-dependent degron (Song et al., 2021 and own observation), we placed the endogenous *dbp2* gene under the control of the *P*.*nmt1* promoter, which is repressed in the presence of thiamine (Siam et al., 2004). We used a Myc-tagged strain to monitor the kinetics of protein depletion after a shift to thiamine-containing medium by Western blot (Figure 4A). Based on the depletion time course and strain growth on rich medium (Figure 4B), we chose a duration of 5h for *P*.*nmt-dbp2* depletion.

**Figure 4:**
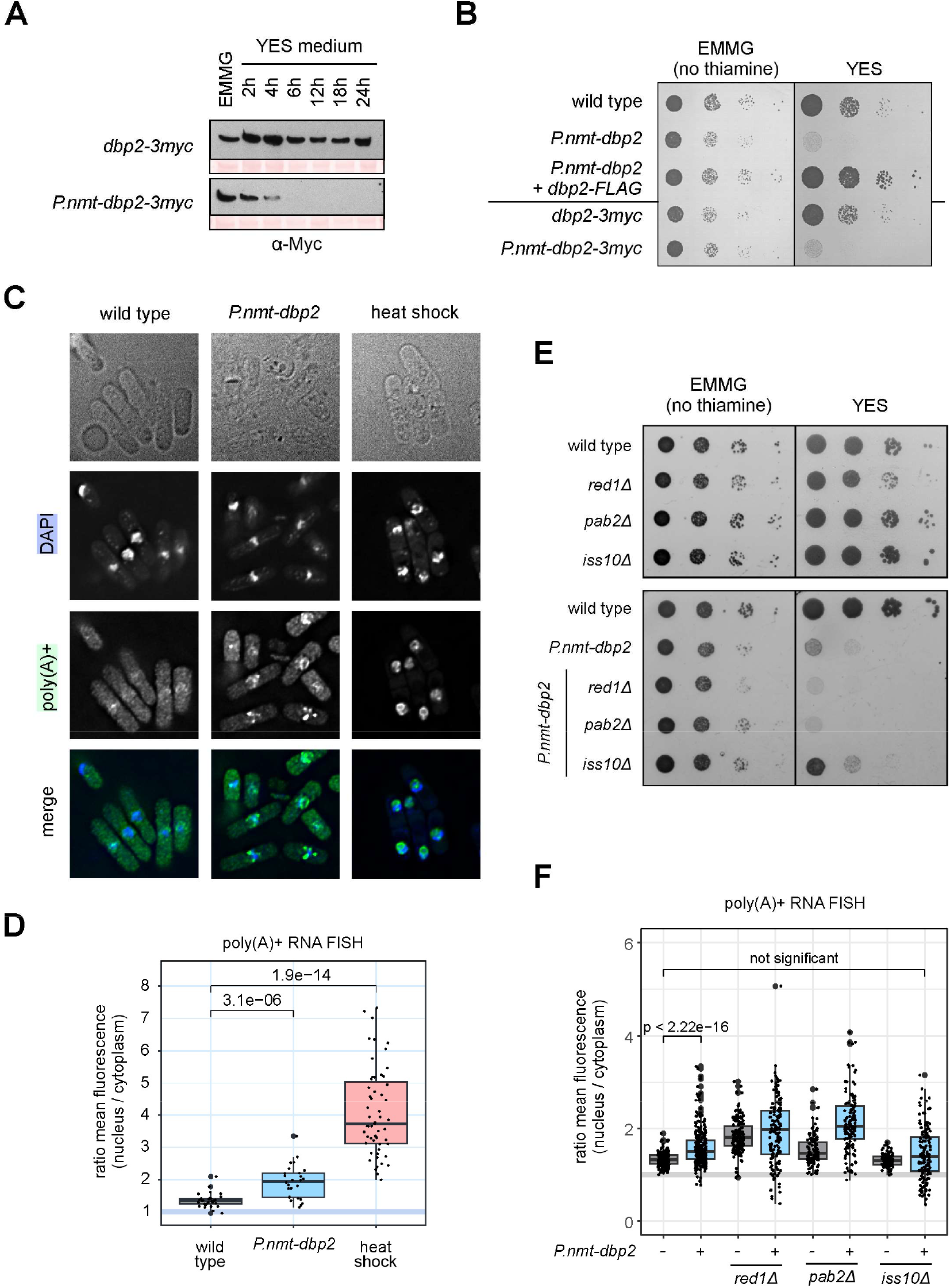
In the absence of Dbp2, 3’-processed RNA accumulates in the nucleus. **A** Western blot depletion time course for Dbp2-3myc under its endogenous or the P.*nmt1* promoter, respectively. Cells were grown in EMMG overnight at 30°C, harvested, and cultured in YES medium for the indicated amount of time. 10 µg protein were loaded per lane and the Myc-tagged protein detected using a mouse monoclonal α-c-Myc antibody, clone 9E10. Ponceau straining of the membrane is included as a loading control. **B** In a plate-based growth assay, metabolic depletion of Dbp2 or Dbp2-3myc under the *P*.*nmt1* promoter leads to poor growth on thiamine-containing medium (YES). The growth phenotype is rescued when an additional copy of *dbp2-FLAG* under its endogenous promoter is inserted at an ectopic locus (*leu1*). The indicated strains were grown in EMMG overnight and serial dilutions (1:10) were spotted on EMMG or YES and incubated at 30°C. Image is representative of three independent experiments. **C** Fluorescence in-situ hybridization (FISH) against poly(A)+ RNA using a Cy3-labelled oligo-d(T) probe and DAPI to stain the DNA within the nucleus. Cells were grown overnight in EMMG, then shifted to YES for 5h to shut off the *P*.*nmt1* promoter before formaldehyde fixation. For the heat shock control, wild-type cells were shifted to 42°C for 1h. Images are representative of two independent experiments. The merged channel shows poly(A)+ RNA in green and DAPI in blue. A quantitation of the nuclear/cytoplasmic signal distribution is given in B. **D** Quantitation of the oligo-d(T) FISH experiment shown in 4C (n = 2). Semi-automated cellular segmentation was carried out by thresholding on the transmitted light channel (cell outlines) and DAPI stain (nuclei). Measurements were performed on average intensity Z-projections for nucleus and cytoplasm (= total cell without nucleus) and the ratio of mean nuclear fluorescence intensity over mean cytoplasmic fluorescence intensity calculated for each cell. The displayed p-values for the pair-wise comparisons were calculated using the Wilcoxon test. Note that the calculated ratios underestimate nuclear RNA retention, particularly in the heat shock sample, because segmentation on the DAPI channel is biased towards the DNA-rich nuclear compartment and can exclude parts of the nucleolus. **E** Growth assay of MTREC mutants combined with *P*.*nmt-dbp2*. The indicated strains were grown in EMMG overnight, and serial dilutions (1:10) were spotted on EMMG or YES and incubated at 30°C (n = 2). Deletion of *iss10* enhances growth in the absence of Dbp2. **F** poly(A)+ RNA FISH experiment of MTREC mutants combined with *P*.*nmt-dbp2* as in C and D (n = 3). At least a hundred cells were counted for each condition. Representative fluorescence images are provided in Suppl. Figure 4C.

To determine the localization of RNAs that have undergone processing by cleavage and polyadenylation, we performed poly(A)+ RNA FISH on formaldehyde-crosslinked cells. As a control, we included a heat shock treatment, which is known to efficiently block bulk mRNA export in yeast (Saavedra et al., 1996). In wild-type cells, poly(A)+ RNA is evenly distributed throughout the cytoplasm and moderately more abundant in the nucleus, with the mean ratio of average fluorescence intensities of nucleus and cytoplasm at 1.36 (Figure 4C and D). Upon heat shock, we observed almost complete nuclear retention of poly(A)+ RNA, which accumulates in a striking ring-shaped pattern from which chromatin appears to be excluded (compare DAPI stain). These poly(A)+ RNA-rich structures are likely to correspond to nucleolar rings, in which essential factors of nuclear RNA metabolism have been shown to reversibly aggregate during heat stress (Gallardo et al., 2020). Upon depletion of Dbp2, poly(A)+ RNA is still present in the cytoplasm; however, we observed a highly significant increase in nuclear to cytoplasmic signal (1.92; p-value = 3.1e-06 (Wilcoxon)). We conclude that 3’-processed, polyadenylated RNAs are inefficiently exported into the cytoplasm in the absence of Dbp2.

Although much of the poly(A)+ RNA that was retained in the nucleus in the *P*.*nmt-dbp2* mutant was dispersed in the nucleoplasm, we noted that the signal partially aggregated in foci. As we had previously determined that Dbp2 localizes to cleavage bodies in wild-type cells, we wondered whether these would be the sites where poly(A)+ RNA accumulated in its absence. Using FISH in *P*.*nmt-dbp2* cells in which the cleavage body component Mtl1 was C-terminally tagged with GFP, we confirmed that the sites of poly(A)+ RNA accumulation correspond to cleavage bodies, suggesting that Dbp2 is important for the release of poly(A)+ RNAs from this compartment (Suppl. Figure 4A and B).

### MTREC/PAXT components Red1 and Pab2 are not required for the nuclear retention of poly(A)+ RNA in the absence of Dbp2

Dbp2 is an essential gene in *S. pombe*. Based on our observations, it is conceivable that the failure to efficiently export poly(A)+ RNA contributes to the poor growth phenotype of the depletion strain. Therefore, we hypothesized that combining a mutation that alleviates RNA retention with Dbp2 depletion could increase viability on a thiamine-containing medium. Recent work in human cells has identified the Red1 homologue ZFC3H1 as a nuclear retention factor for exosome target RNAs that can sequester RNA in nuclear condensates in a manner dependent on the nuclear poly(A)-binding protein PABPN1 (Fan et al., 2018; E. S. Lee et al., 2022; Silla et al., 2018; Y. Wang et al., 2021). To test the involvement of MTREC/PAXT in poly(A)+ RNA retention in the *dbp2* mutant, we crossed *red1Δ, pab2Δ* (homologue of PABPN1) and *iss10Δ* into the *P*.*nmt-dbp2* background. Iss10 is a fission yeast-specific component of MTREC, which shows high homology to the N-terminal region of ZFC3H1 and is required for cleavage body integrity (Egan et al., 2014; Yamashita et al., 2013; Zhou et al., 2015). Combining *P*.*nmt-dbp2* with *red1Δ* or *pab2Δ* did not improve growth on YES; rather, it made it slightly worse (Figure 4E). In contrast, combination with *iss10Δ* provided *P*.*nmt-dbp2* with a mild but reproducible growth advantage under depletion conditions (Figure 4E).

To determine the impact of the additional mutations on poly(A)+ RNA export, we carried out poly(A)+ RNA FISH experiments. The single deletion of *red1Δ* or *pab2Δ* led to a retention of poly(A)+ RNA in foci even in the presence of Dbp2, with an additive effect on export if both mutations were combined (Figure 4F and Suppl. Figure 4C). The single *iss10Δ* deletion, on the other hand, was undistinguishable from the wild type. Combining *iss10Δ* with *P*.*nmt-dbp2* reduced nuclear poly(A)+ RNA retention compared to *P*.*nmt-dbp2* alone, and the ratio of average fluorescence intensities of nucleus and cytoplasm was no longer statistically significantly different from the wild type; however, it should be noted that *P*.*nmt-dbp2* displays a much larger cell-to-cell variability of observed nuclear over cytoplasmic signal ratios in any genetic context, including in combination with *iss10Δ* (Figure 4F), and that cells with strong punctate poly(A)+ RNA signal in the nucleus are still frequently observed in the *iss10Δ P*.*nmt-dbp2* double mutant. We conclude that additional, unidentified factors contribute to RNA retention in the *P*.*nmt-dbp2* background.

### CPAC components are depleted from the soluble pool upon loss of Dbp2

If Dbp2 played an active role in CPAC release in the context of an mRNP assembly checkpoint, its depletion would be expected to lead to increased CPAC retention. To determine whether Dbp2 was involved in CPAC recycling, we first examined the subcellular distribution of CFIA component Pcf11 in the presence and absence of Dbp2. In wild-type cells, the majority of GFP-tagged Pcf11 colocalized with Red1-tdTomato in cleavage bodies, with little fluorescence detectable in the nucleoplasm (Figure 5A, upper panel, and 5B, left panel). Upon depletion of Dbp2, we observed a distinct redistribution of Pcf11-GFP: Colocalization with cleavage bodies was markedly reduced, and much of the fluorescent signal was now dispersed throughout the nucleoplasm (Figure 5A, lower panel, and 5B, right panel). Based on our earlier results, we assume that this reflects a redistribution from the storage compartment to the sites of RNA biogenesis.

**Figure 5:**
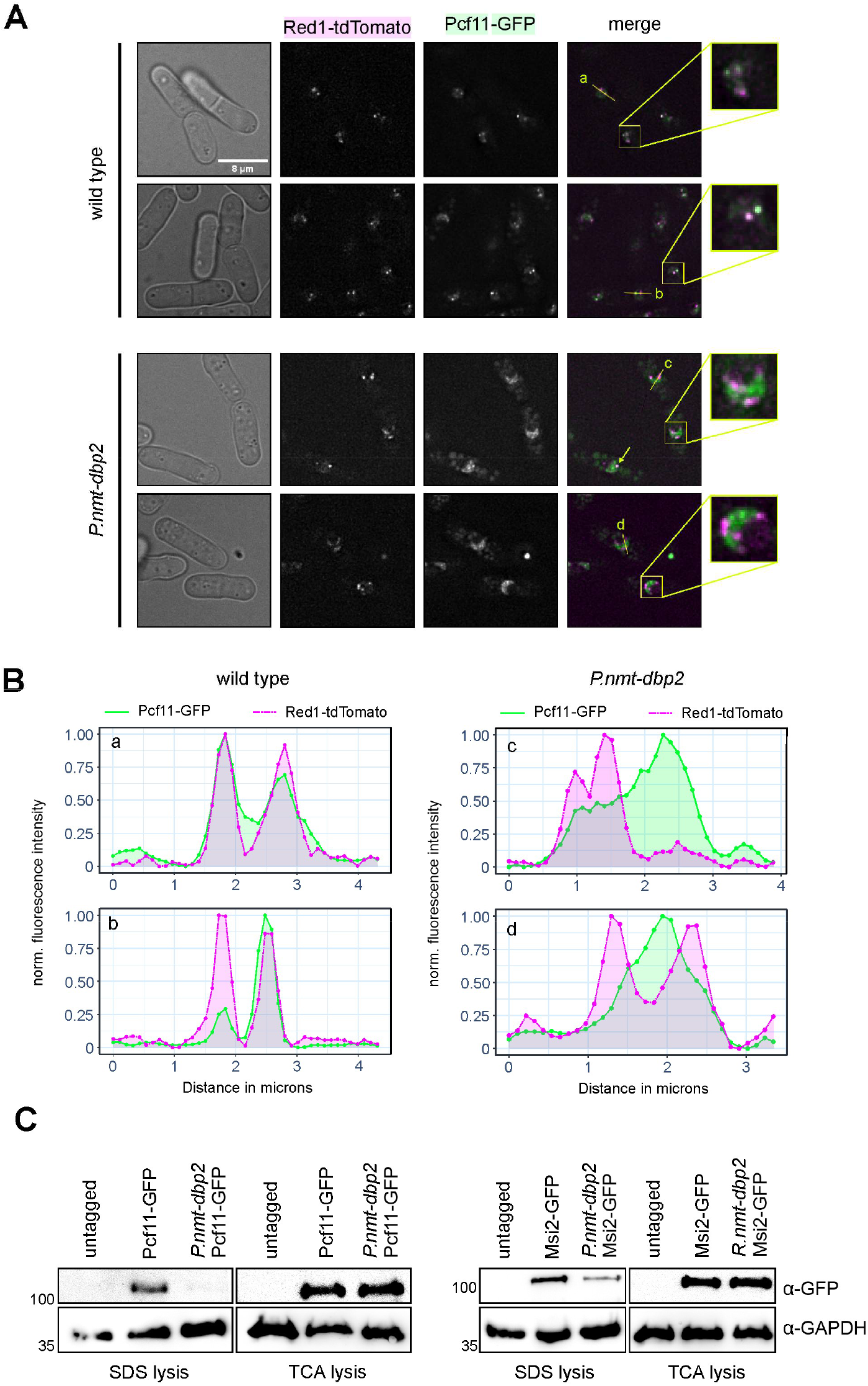
CPAC components are depleted from the soluble pool in the absence of Dbp2. **A** Live cell imaging of genomically tagged Pcf11-GFP and the MTREC component Red1-tdTomato. Cells were grown in YES at 30°C, pelleted and resuspended in EMMG for imaging on poly-lysine-coated coverslips. The merged channel shows Pcf11-GFP in green and Red1-tdTomato in magenta. Images are representative of three independent experiments. Fluorescence intensity profiles were generated along the yellow lines and are shown in B. Enlarged versions of the merged image without the yellow markings are provided in Suppl. Fig. 5A. **B** Fluorescence intensity profiles across cleavage bodies and the adjacent nuclear area as indicated in A. Green line corresponds to the Pcf11-GFP signal, dashed magenta line to the Red1-tdTomato signal. Fluorescence intensities were measured along the line and normalised to a 0-1 range. **C** Western blot analysis of cell extracts generated by SDS lysis (soluble cell extracts) or TCA lysis (total cell extracts) for CFIA component Pcf11-GFP (left panels) and CFIB component Msi2-GFP (right panels). Cells were grown in EMMG overnight at 30°C, harvested, and cultured in YES for 5h before lysis. GAPDH was included as a loading control. The numbers on the left indicate the molecular weight marker in kDa. Images are representative of two independent experiments.

We had noted earlier that lysates prepared from the *P*.*nmt-dbp2* strain appeared to contain much lower levels of various CPAC components than the wild type. However, this disagreed with fluorescence microscopy data, where we observed no change in the total fluorescent signal of GFP-tagged CPAC components between wild type and mutant (compare Figure 5A). It has been described previously that stalling of RNP complexes on chromatin can interfere with their efficient extraction using standard protocols for chromatin preparation (Rougemaille et al., 2008). We wondered whether a similar effect led to the apparent loss of CPAC components from lysates in the *dbp2* mutant. To test this hypothesis, we compared protein amounts in standard SDS lysates with extracts prepared using the TCA method, which has been reported to be much more efficient in extracting chromatin (Mehta et al., 2010). Indeed, while Dbp2 depletion strongly reduced the amounts of Pcf11 or of the CFIB component Msi2 recovered after SDS lysis, their yield after TCA lysis was not affected (Figure 5C). This suggests that CPAC components are depleted from the soluble pool in the absence of Dbp2, possibly because they are sequestered on poly(A)+ RNA retained in the nucleus and/or on chromatin.

### Dbp2 depletion increases skipping of cleavage and polyadenylation sites

Assuming that CPAC components in the soluble pool correspond to the idle fraction that is available for recruitment to transcribing RNAPII for the 3’-end processing of nascent transcripts, we wondered whether reduced levels of soluble CPAC in the *dbp2* mutant would lead to inefficient cleavage and polyadenylation. We performed RNA-seq analysis on poly(A)-selected RNA to determine whether depletion of Dbp2 had an impact on PAS usage. We used *S. cerevisiae* cells as a spike-in for normalization to be able to assess the global impact on mRNA expression levels. We observed a transcriptome-wide reduction in RNA levels after Dbp2 depletion, in agreement with an essential function for Dbp2 as a global regulator of RNAPII-dependent RNA biogenesis (Figure 6A and Suppl. Figure 6A). Similar results were obtained when we sequenced ribodepleted RNA after a 9h depletion period (Suppl. Figure 6A). Because Dbp2 co-purified and co-localized with MTREC/PAXT, we also verified whether decay of exosome substrates was affected. Levels of known MTREC-dependent RNA targets of the nuclear exosome, including meiotic mRNAs that are part of the Mmi1 regulon and snoRNA precursors that are dependent on the nuclear membrane protein Lem2 (Chen et al., 2011; Kilchert et al., 2015; Martín Caballero et al., 2022), were not increased relative to other transcripts upon depletion of Dbp2 (Figure 6B), indicating that Dbp2 is not required for MTREC-dependent RNA turnover.

**Figure 6:**
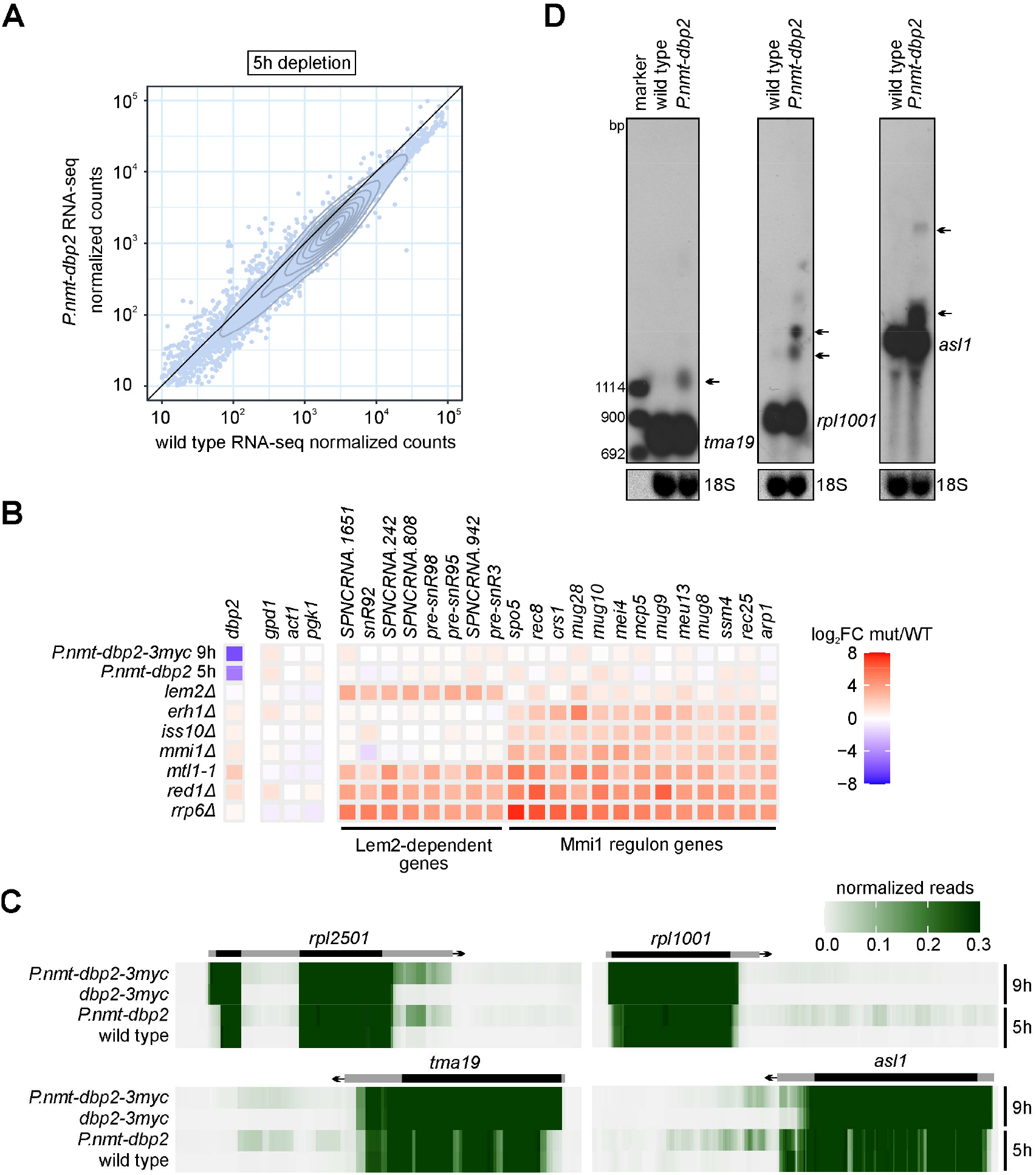
Dbp2 depletion increases skipping of cleavage and polyadenylation sites. **A** Mean integrated counts of RNA-seq reads over annotated features in wild-type and *P*.*nmt-dbp2* (n = 3) after a 5h depletion period. Cells were grown in EMMG overnight at 30°C, harvested, and cultured in YES for 5h, then mixed with *S. cerevisiae* in a 5:1 OD_600_ ratio prior to RNA isolation. Integrated counts were normalized to the number of total *S. cerevisiae* reads for each sample before calculating the mean. Contours of a 2d density estimate are shown in dark grey. **B** DESeq2 differential expression analysis of nuclear exosome target genes that are either Lem2-dependent or part of the Mmi1 regulon (Chen et al., 2011; Martín Caballero et al., 2022); *adh1, pgk1* and *act1* are housekeeping mRNAs and included as controls. Colour indicates log_2_ fold change of normalized RNA-seq counts of mutant over wild type. MTREC mutant data from (Atkinson et al., 2018) (*rrp6Δ*), (Kilchert et al., 2015) (*mmi1Δ*), (Birot et al., 2021) (*mtl1-1*), and (Martín Caballero et al., 2022) (*red1Δ, iss10Δ, erh1Δ, lem2Δ)*; Accessions PRJEB7403, GSE73144, GSE148799, and GSE174347. *S. cerevisiae* spike-in was disregarded for the analysis to obtain comparable relative expression data. **C** Heat maps of RNA-seq reads across *rpl2501, tma19, rpl1001*, and *asl1*, and downstream regions in untagged (n = 3) or 3myc-tagged (n = 2) Dbp2 after a 5h or 9h depletion period, respectively. Reads were counted across 10 bp bins, and the means normalized to a 0-1 range and plotted at y_max_ = 0.3 to visualize relative amounts of 3’-extended transcripts. The positions of the annotated transcripts are indicated in grey, the coding sequence is highlighted in black. **D** Northern blot for *tma19, rpl1001*, and *asl1* mRNA using a strand-specific DIG-labelled RNA probe against the gene body. Cells were grown in EMMG overnight at 30°C, harvested, and cultured in YES medium for 5h prior to RNA isolation. 18S band stained with methylene blue is shown as a loading control. Arrows indicate positions of extended transcripts. The marker is a double-stranded DNA size marker (Roche, Molecular Weight Marker VIII, DIG-labeled).

To determine the efficiency of cleavage and adenylation, we assessed PAS usage. For many mRNAs, we observed a relative increase in sequencing reads over longer 3’-UTR isoforms or past the CPA site of annotated features after Dbp2 depletion, revealing a general tendency of the mutant to skip PAS’s and continue transcription to downstream sites (Figure 6C and Suppl. Figure 6B and C). Northern blotting with probes directed against the gene bodies confirmed the presence of extended transcript isoforms for several mRNAs that we tested (Figure 6D and Suppl. Figure 6D). In the RNA-seq data, increased read-through signal could be detected for almost any transcript with a sufficiently high expression level; however, the severity of the phenotype was transcript-dependent and may be related to the strength of the corresponding PAS.

### In the absence of Dbp2, transcription termination is delayed

Because transcription termination is coupled to co-transcriptional RNA cleavage, we also sought to directly determine the impact of Dbp2 depletion on RNAPII transcription. Using spike-in controlled RNAPII-ChIP-seq, we observed a global reduction of RNAPII levels at transcribed genes upon depletion of Dbp2 (Figure 7A and B). To determine whether transcription termination was affected, we performed metagene analysis of RNAPII coverage at RPGs. In the absence of Dbp2, the peak in RNAPII coverage at the 3’-end of genes was shifted downstream compared to the wild type, indicating a delay in transcription termination (Figure 7C, left panel). According to the prevalent model of cleavage-coupled transcription termination, RNAPII is in kinetic competition with the 5’-3’ exonuclease Dhp1 (homologue of Xrn2), which degrades the nascent 3’-fragment that is generated during cleavage and promotes transcription termination once it reaches RNAPII (Fong et al., 2009; M. Kim et al., 2004; Larochelle et al., 2018; West et al., 2004). Because of this competition, both faster RNAPII elongation rates and a delay in RNA cleavage can delay transcription termination. At present, we cannot unambiguously distinguish between these possibilities. However, relative to the amount of RNAPII that reaches the PAS, the height of the RNAPII peak across the termination zone is increased upon Dbp2 depletion, which is indicative of a slow transition of RNAPII through the area (Figure 7C, right panel). We therefore consider it likely that depletion of Dbp2 decreases the efficiency of CPA at RNAPII-transcribed genes, which is consistent with the increased occurrence of PAS skipping that we observed on the RNA level. We take this as an indication that the depletion of CPAC components from the soluble pool in the absence of Dbp2 limits their availability to levels that are insufficient to maintain normal levels of 3’-end processing.

**Figure 7:**
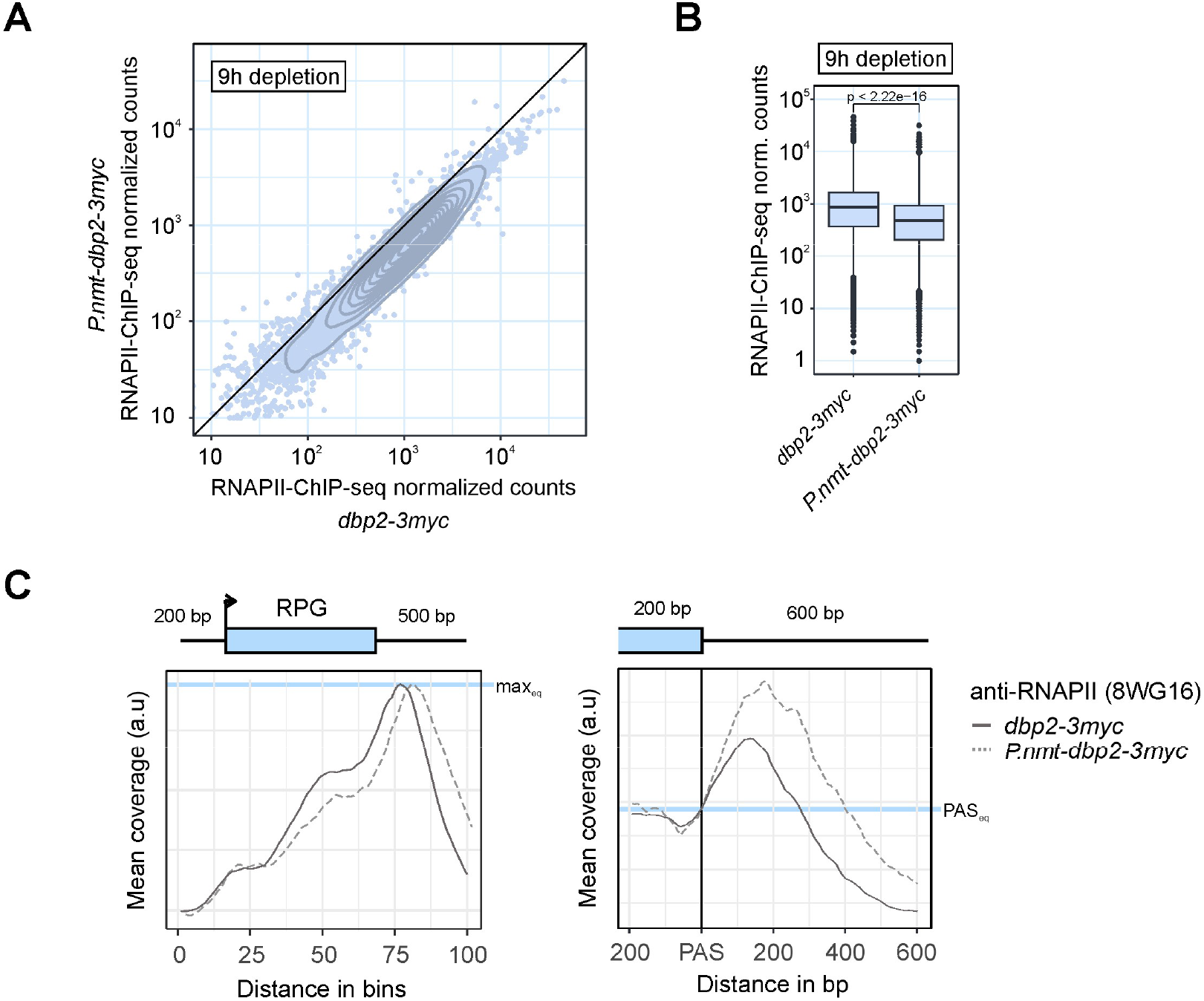
In the absence of Dbp2, transcription termination is delayed. **A** Mean integrated counts of RNAPII-ChIP-seq reads over annotated features in d*bp2-3myc* or *P*.*nmt-dbp2-3myc* (n = 2) after a 9h depletion period. Cells were grown in EMMG overnight at 30°C, harvested, and cultured in YES for 9h, then mixed with *S. cerevisiae* in a 5:1 OD_600_ ratio prior to chromatin isolation and IP against total RNAPII (α-rpb1 (8WG16), n=2). Integrated counts were normalized to the number of total *S. cerevisiae* reads for each sample before calculating the mean. Contours of a 2d density estimate are shown in dark grey. **B** Mean integrated counts of RNAPII-ChIP-seq reads over annotated features as in A. The displayed p-value for the pair-wise comparison was calculated using the Wilcoxon test. **C** Metagene analysis of mean RNAPII ChIP-seq signal (n = 2) across ribosomal protein genes (RPGs) including 200 bp upstream and 500 bp downstream of the annotated transcription units (left panel) or surrounding the polyadenylation and cleavage site (PAS) (right panel). Mean coverage was adjusted by a constant scaling factor to normalize to maximal peak height (left panel) or RNAPII levels at the PAS (right panel) for an easier comparison of curve shapes. Schematic of the gene above the left panel corresponds to an RPG of median length. In the left panel, the positions of transcription start and end sites within the metagene are distributed around the given coordinate because of the varying feature compression depending on gene length.

## Discussion

Despite an extensive body of work that has advanced our understanding of the biological function of the RNP remodelling helicase Dbp2/DDX5, the mechanisms underlying the pleiotropic phenotypes associated with its dysfunction are still elusive. In this study, we provide evidence that places fission yeast Dbp2 at the intersection between RNA 3’-end formation and RNA export. Our data is consistent with a model in which Dbp2 is the key enzyme in an early mRNP remodelling checkpoint that is coupled to the release of CPAC into the soluble pool and allows 3’-end processed transcripts to gain export competence. Importantly, this would establish transcript release after 3’-end formation as an ATP-dependent process.

The existence of such a checkpoint has been proposed based on the analysis of RNA export mutants in *S. cerevisiae* (Qu et al., 2009). Here, temperature-sensitive alleles of the export receptor Mex67, the export adaptor Yra1, and various other RNA assembly factors led to the retention of the CFIA components Rna14 and Rna15 on polyadenylated RNA. Conversely, some mutations in CFIA that retain the ability to support cleavage and polyadenylation lead to a failure to export the processed transcripts (Hammell et al., 2002). Notably, our comparative interaction profiling experiment found several RNA export factors to be enriched in the Dbp2-HTP purification, among them Mex67 and the Yra1 homologue Mlo3; in *S. cerevisiae, Sc*Mex67 and Yra1 have also been shown to interact with *Sc*Dbp2 and regulate its function, and deletion of *DBP2* – which is not an essential gene in budding yeast – was shown to reduce association of Mex67 with transcripts (Ma et al., 2013, 2016). In *Drosophila*, a strong similarity between the phenotypes of mutants of the Mex67 and Dbp2 homologues *small bristles* and *Rm62* has been described (Buszczak & Spradling, 2006). One hypothesis would be that direct interaction between export factors and Dbp2 initiates CPAC displacement. It will therefore be of great interest to investigate the mode of interaction between Dbp2 and these export factors in further studies.

At present, we can only speculate about the mechanism of RNA retention in the absence of Dbp2. A non-negligible amount of the nuclear poly(A)+ RNA signal in *P*.*nmt-dbp2* is present in cleavage bodies (Suppl. Figure 4A and B), suggesting that they might serve a function analogous to nuclear speckles (NS) in metazoans. NS are enriched in splicing factors and have been proposed to either serve as an active buffering compartment for splicing factors or to represent processing hubs (Hirose et al., 2022). They are also the sites where poly(A)+ RNA accumulates upon an export block; therefore, passage through NS is thought to allow mRNAs to acquire export competence (K. Wang et al., 2018). However, in contrast to *red1Δ* and *pab2Δ*, where nuclear-retained poly(A)+ RNAs almost exclusively localize to cleavage bodies, disperse poly(A)+ RNA constitutes a significant fraction of the total nuclear signal after Dbp2 depletion (Figure 4C and F). At present, our best assumption is that RNA retention already occurs at an early stage of RNA biogenesis and may represent a failure to release RNA from chromatin after transcription. This interpretation is supported by live cell imaging of CPAC components and their depletion from the readily extractable soluble pool (Figure 5A and C). In murine cells expressing a version of Rpb1 with a truncated CTD, fully processed mRNAs are retained at the transcribed gene, suggesting that release from chromatin constitutes an independent step of gene expression that requires an active component usually recruited by the full-length CTD (Custódio et al., 2007). Recent experiments that measured RNA residence times in various compartments in mammalian cells identified release from chromatin as the rate-limiting step for nuclear export (Smalec et al., 2022); a similar observation had been made earlier for a subclass of mRNAs encoding inflammatory proteins, and in that case had been linked to slow splicing kinetics (Pandya-Jones et al., 2013). Poor RNA processing has long been known to result in the retention of RNAs at the site of transcription, which is supported by genome-wide correlation studies (Custódio et al., 1999; Henfrey et al., 2023; Hilleren et al., 2001; Rougemaille et al., 2008). Whether this retention is mechanistically related to the RNA retention we observe after the loss of Dbp2 is currently unclear. However, in *S. cerevisiae*, the CFIB component and Msi2 homologue Hrp1 was recently shown to be required to prevent the export of incorrectly processed transcripts to the cytoplasm (J. Li et al., 2023). There is relatively little published work that has directly addressed the question of how selective tethering of RNA to chromatin can be achieved: For *S. cerevisiae* mutants of the TREX complex, chromatin retention has been convincingly linked to the formation of R-loops (Penzo et al., 2023). In that case, retention coincides with downregulation of the CPAC component Fip1, a co-factor of the poly(A) polymerase, and defects in polyadenylation (Saguez et al., 2008). Whether the failure to polyadenylate the transcript is a prerequisite for strand invasion and efficient RNA:DNA hybrid formation in TREX mutants is an open question – if it was, it would be unlikely that the poly(A)+ RNA that we observe in the absence of Dbp2 is tethered to chromatin by the same mechanism. Moreover, our protein interaction profiling data do not provide compelling evidence that Dbp2 in *S. pombe* preferentially associates with R-loops under normal growth conditions. As an alternative to direct tethering to DNA as part of an RNA:DNA hybrid, retention on chromatin could also be linked to the formation of RNA-rich protein condensates. In addition to Dbp2 itself, which has the ability to induce phase separation (Hondele et al., 2019), at least one 3’-end processing factor in *S. pombe*, the CTD-interacting protein Seb1 (Lemay et al., 2016; Wittmann et al., 2017), has been described to form chromosome-tethered condensates under specific conditions in a recent preprint (Ding et al., 2023). Alternative mechanisms of RNA retention are conceivable, and contributing factors and the exact mechanism will have to be determined in the future.

### Links to the nuclear RNA surveillance machinery

Many processing defects that induce the retention of specific transcripts on chromatin are linked to the degradation of the retained transcript by the nuclear exosome (Assenholt et al., 2008; Hilleren et al., 2001; Kilchert et al., 2016). Notably, we find that Dbp2 both co-purifies and co-localizes with MTREC, the nuclear exosome targeting complex related to human PAXT composed of the Mtr4-like helicase Mtl1, the zinc-finger protein Red1, and various associated factors including the nuclear poly(A)-binding protein Pab2 (Egan et al., 2014; Kilchert, 2020; N. N. Lee et al., 2013; Meola et al., 2016; Silla et al., 2020; Zhou et al., 2015). To date, we have found no evidence for a direct link between Dbp2 and MTREC function. Dbp2 is not required for the turnover of known exosome targets (Figure 6B), and RNA retention after Dbp2 depletion is not dependent on Red1 or Pab2 (Figure 4F and Suppl. Figure 4C). Rather, the effects of both mutations on RNA retention appear to be additive, suggesting that non-overlapping RNA populations may be affected; this would be consistent with data from human PAXT mutants, in which nuclear RNA accumulations have been shown to harbour RNAs that would usually be degraded by the exosome complex (Silla et al., 2018; Y. Wang et al., 2021). In contrast, we observe a positive genetic interaction between *P*.*nmt-dbp2* and *iss10Δ*, and a mild but significant improvement of RNA export in the double mutant (Figure 4E and F); the molecular basis of this remains unclear. Interestingly, depletion of the exosome complex also causes read-through transcription and delayed transcription termination (Lemay et al., 2014). Metagene analysis of published RNAPII ChIP-seq data for *P*.*nmt-dis3* over the same set of genes reveals very similar changes in RNAPII occupancy as we have observed after Dbp2 depletion (Figure 7C and Suppl. Figure 7A) (Lemay et al., 2014). Moreover, in a comparative RNA interactome capture experiment in which we had previously compared occupancies of RNA-binding proteins on polyadenylated RNA between wild type and *mtl1-1* or *rrp6Δ* (mutants of MTREC and the exosome complex, respectively), CPAC components – in particular CFIA – were significantly enriched, suggesting that CPAC release might be impaired in exosome mutants (Birot et al., 2021; Kilchert et al., 2019) (Suppl. Figure 7B and C). However, while the strong similarities between the phenotypes for *dbp2* and exosome mutants are suggestive, further studies are required to develop a robust theory of how the two are mechanistically linked.

### Comparative interaction profiling reveals sequence of co-transcriptional events

While the chromatin association of both Srp2 and Dbp2 covers the entire transcription unit of RNAPII-transcribed genes, our data suggest that Srp2 associates with transcribing RNAPII earlier during the transcription cycle than Dbp2 and is also released earlier (Figure 1C). Consequently, the relative enrichment scores of proteins in the comparative protein interaction profiling data of Srp2-HTP and Dbp2-HTP appear to reflect how early or late a factor is recruited to the RNAPII holocomplex during the transcription cycle. In general, subunits of the same complex tend to have very similar enrichment scores, which greatly adds to the confidence with which we can assign scores to complexes. This proposed relationship between relative enrichment and time of arrival holds for many factors for which the timing of recruitment is known (Figure 2D and E). It is therefore tempting to speculate that this dataset may also have predictive value for factors for which the order of recruitment is currently unclear. For example, the strong enrichment of TREX in the Srp2-HTP purification (Figure 2E and Suppl. Figure 2D) suggests that the mode of TREX recruitment in *S. pombe* may reflect the metazoan situation, where TREX is recruited to the 5’-end of genes in a splicing-dependent manner, in contrast to *S. cerevisiae*, where TREX is recruited by a transcription-coupled mechanism (Cheng et al., 2006; Masuda et al., 2005; Zenklusen et al., 2002). It is unclear to what extent the exon junction complex (EJC) in *S. pombe* resembles the mammalian complex and whether it can interact with TREX in a manner similar to the human complex (Marayati et al., 2016; Pacheco-Fiallos et al., 2023); however, its relative enrichment score is similar to that of TREX, suggesting a possible interdependence in recruitment (Figure 2E and Suppl. Figure 2D). To add another example, our data would place the arrival of the Paf1 complex after the recruitment of P-TEFb but ahead of CTDK; this is consistent with published data that show a requirement of P-TEFb activity for the efficient recruitment of the Paf1 complex (Liu et al., 2009; Mbogning et al., 2013; Wier et al., 2013). At the time, we are lacking high-resolution ChIP-seq data for these factors to validate these predictions. Nevertheless, we suggest that comparative interactome profiling of factors associated with different stages of transcription constitutes a valid experimental approach to determine the sequence of co-transcriptional events.

### ATP-dependence of RNA-protein complex dissolution as a general principle in RNA metabolism

Finally, we want to point out that in the splicing field, the importance of DEAD- and DEAH-box ATPases in reversing the highly specific interactions that underlie splice site recognition and stabilize the various conformations of the spliceosome – thus enabling the highly dynamic, multi-step splicing reaction to proceed – has long been recognized (Enders et al., 2023). Viewed in this context, it may come as no surprise that the release of CPAC is coupled to ATP hydrolysis via the action of a DEAD-box ATPase, Dbp2. Whether energy expenditure for complex dissolution would be the price to be paid for the high selectivity of the initial RNA-protein interaction or simply a means to introduce an additional layer of regulation remains a matter of speculation.

## Methods

### Yeast Strains and Manipulations

All *S. pombe* strains used in this study are listed in Table S2. Standard methods were used for cell growth and genetic manipulations (Bähler et al., 1998; Knop et al., 1999; Matsuyama et al., 2004; Moreno et al., 1991; Zhang et al., 2022). Oligos and plasmids used for strain construction are listed in Tables S3 and S4. Sequences and functional annotations were retrieved from PomBase (Harris et al., 2022). Cells were grown in yeast extract with supplements (YES) or Edinburgh minimal medium with glutamate (EMMG) at 30°C unless indicated otherwise.

### Chromatin immunoprecipitation

ChIP-seq was carried out as described (Wittmann et al., 2017), with minor modifications. Exponentially growing cells (200 ml) were crosslinked with 1% formaldehyde for 20 min at room temperature (RT). For calibrated ChIP-seq of RNAPII, *S. cerevisiae* cells were added to the culture in a 1:5 OD_600_ ratio prior to crosslinking. 30 ml of a solution of 3 M glycine, 20 mM Tris were added to quench the reaction. Cells were pelleted and washed once with cold TBS and once with FA lysis buffer / 0.1% SDS (50 mM Hepes-KOH pH 7.5, 150 mM NaCl, 1 mM EDTA, 1% Triton0020X-100, 0.1% sodium deoxycholate). After resuspension in 1.4 ml FA lysis buffer / 0.5% SDS, cells were distributed to two screw-cap microtubes containing 300 µl glass beads and lysed in a FastPrep instrument (MP Biomedicals) at 3 x 45 s 6 m/s, with 2 min breaks on ice. Lysed cells were recovered with FA lysis buffer / 0.1% SDS and ultracentrifuged (24,000 rpm, 20 min, 4°C) in a 70.1 Ti rotor. The chromatin pellet was resuspended in FA lysis buffer / 0.1% SDS and sheared with a Bioruptor sonicator (Diagenode) at 15 s ON / 45 s OFF for 80 min. HTP-tagged proteins were immunoprecipitated with IgG-coupled Dynabeads (tosylactivated M280, Thermo Scientific), RNAPII with antibody against Rpb1 (8WG16, Millipore) coupled to 20 µl of protein-G dynabeads (Life Technologies). After washing and elution of bound material from the beads, proteins were digested by incubation with 0.6 mg pronase for 1 h at 42 °C, followed by decrosslinking at 65 °C overnight and DNA extraction. Sequencing libraries were constructed with the TruSeq ChIP Library Preparation Kit (Illumina) and sequenced on a NextSeq 2000 instrument (Srp2-HTP & Dbp2-HTP) or with the NEBNext Fast DNA Library Prep Set for Ion Torrent Kit (NEB) and sequenced on an Ion Torrent Proton (Life Technologies) (RNAPII-ChIP).

### Comparative protein interactome profiling

Cells were grown in 2 l YES to an OD_600_ of 3.9 and crosslinked with 0.01% formaldehyde for 10 minutes at 30°C before harvesting. Cell pellets were washed once with lysis buffer (20 mM HEPES pH 8.0, 100 mM KAc, 2 mM MgCl_2_, 3 mM EDTA, 0.1% NP-40, 10% glycerol, 1 mM DTT, protease inhibitor (P8215, Sigma)) and snap frozen in liquid nitrogen until use. Cells were lysed by grinding under liquid nitrogen for 30 min, and lysates pre-cleared at 4,000 rpm for 12 minutes at 4°C (3488 rcf). Lysates were cleared by centrifugation for 1 h at 45,000 rpm at 4°C (Optima XPN-80 Ultracentrifuge with 70 Ti, Beckman Coulter). HTP-tagged proteins were immunoprecipitated with 600 µl pre-washed IgG-sepharose 6 fast flow affinity resin (GE Healthcare) for 2 hours at 4°C in a rotating wheel. After washing, proteins were eluted by TEV cleavage (20 µg TEV in 200 µl lysis buffer for 75 min at 16°C) with 20 µg home-made TEV enzyme. The TEV eluate was incubated with 200 µl pre-washed Protino® Ni-NTA Agarose beads (Macherey-Nagel) for 1 h at 4°C in a rotating wheel. After washing (20 mM HEPES pH 8.0, 250 mM NaCl, 10 mM imidazole), purified proteins were eluted in elution buffer (20 mM HEPES pH 8.0, 150 mM NaCl, 300 mM imidazole) for 30 minutes at 4°C in a rotating wheel.

### Detection of protein interactor by liquid chromatography-mass spectrometry

The eluate was then precipitated using 7x volumes acetone for at least 2h at -20 °C. Following centrifugation, the protein pellet was washed twice with 300 µl ice cold acetone. The protein pellet was then dried and reconstituted in 100 mM ammonium bicarbonate containing 1 mM TCEP and incubated at 90 °C for 10 min. Alkylation of reduced disulfide bonds was performed with 5 mM iodoacetamide for 30 min at 25°C in the dark. Proteins were digested with 1 µg trypsin at 30°C overnight. The sample was then acidified using trifluoroacetic acid and peptides were purified using Chromabond C18 Microspin columns (Macherey-Nagel).

Dried peptides from on-bead digests were reconstituted in 0.1% trifluoroacetic acid and then analyzed using liquid chromatography mass spectrometry carried out on a Q-Exactive Plus instrument connected to an Ultimate 3000 RSLC nano and a nanospray flex ion source (all Thermo Scientific). Peptide separation was performed on a reverse phase HPLC column (75 μm x 42 cm) packed in-house with C18 resin (2.4 μm; Dr. Maisch). The following separating gradient was used: 98% solvent A (0.15% formic acid) and 2% solvent B (99.85% acetonitrile, 0.15% formic acid) to 35% solvent B over 36 minutes at a flow rate of 300 nl / min. The data acquisition mode was set to obtain one high resolution MS scan at a resolution of 70,000 full width at half maximum (at *m*/*z* 200) followed by MS/MS scans of the 10 most intense ions. To increase the efficiency of MS/MS attempts, the charged state screening modus was enabled to exclude unassigned and singly charged ions. The dynamic exclusion duration was set to 30 sec. The ion accumulation time was set to 50 ms (MS) and 50 ms at 17,500 resolution (MS/MS). The automatic gain control (AGC) was set to 3x10^6^ for MS survey scan and 1x10^5^ for MS/MS scans.

Due to an instrument upgrade co-IP peptide samples generated as described under (2)m, were analyzed on an Exploris 480 (Thermo Scientific) with identical LC settings. The gradient was adjusted to separate peptides with 2 % solvent B to 35 % solvent B over 30 min. MS settings were set as follows: Spray voltage 2.3 kV, ion transfer tube temperature 275 °C, the mass of m/z 445.12003 was used as internal calibrant, MS1 resolution was set to 60.000 with max ion injection time of 25 ms and an automatic gain control (AGC) of 300 %. The RF lens was 40 %. MS/MS scans (cycle 1 s) were carried out with an Orbitrap resolution of 15.000 with an ACG setting of 200 %, quadrupole isolation was 1.5 m/z, collision was induced with an HCD collision energy of 27 %.

MS raw data was then analyzed with MaxQuant (Version 2.0.3.0) (Cox & Mann, 2008), and a *S. pombe* uniprot database. MaxQuant was executed in standard settings without “match between runs” option. The search criteria were set as follows: full tryptic specificity was required (cleavage after lysine or arginine residues); two missed cleavages were allowed; carbamidomethylation (C) was set as fixed modification; oxidation (M), deamidation (N,Q), dimethyl (KR), phospho (STY) as variable modifications. The MaxQuant “proteinGroup.txt” file was further processed with SafeQuant (Ahrné et al., 2013; Glatter et al., 2012) to apply moderated t-statistics for p-value calculation (Kammers et al., 2015) on MaxQuant “LFQ values”.

### Immunofluorescence

Immunofluorescence in Mtl1-GFP was carried out essentially as described (Hagan, 2016). In short, 9 ml cell culture were fixed by the addition of 1 ml 37% formaldehyde for 30 min at RT on a rotating wheel. Cells were washed three times with PEM (100 mM piperazine-N,N′-bis(2-ethanesulfonic acid) (PIPES), 1 mM EGTA pH 8.0, 1 mM MgSO_4_, pH 6.9) and spheroplasted in 500 µl PEMS (PEM with 1.2 M sorbitol) with 12.5 µl 10 mg/ml zymolyase T100 (Carl Roth) for 1h at 37°C. Fixed cells were permeabilised in 500 µl PEMS, 1% Triton X for 30 s, washed three times with PEM, then rotated for 30 min in 200 µl PEMBAL (PEM with 1% bovine serum albumin, 100 mM lysine hydrochloride, 0.1% NaN_3_) at RT. 1/5^th^ of the total fixed cell pellet was used per condition. Cells were incubated in 100 µl PEMBAL with anti-RNA polymerase II CTD repeat YSPTSPS (phospho S2) (Abcam, ab252855, 1:1000) at 4°C overnight. Cells were washed twice with PEMBAL and incubated in 200 µl PEMBAL for 30 min at RT. Cells were incubated in 100 µl PEMBAL with Alexa Fluor 594-conjugated AffiniPure Goat Anti-Rat (Biozol, JIM-112-585-003, 1:400) at 4°C overnight in the dark. Cells were washed once with PEMBAL and once with PBS, then resuspended in 40 µl PBS. 3 µl cell suspension were freshly mounted on poly-lysine-coated coverslips, let to dry, and imaged under ROTIMount FluorCare DAPI (Carl Roth).

### Fluorescence microscopy

Z-stack images were acquired on a Deltavision Ultra High-Resolution Microscope (Cytiva) and deconvolved with the softWoRx software using default settings. Line profiles were generated with Fiji (Schindelin et al., 2012) and plotted in R. For live-cell imaging, cells were grown overnight, harvested, and mounted on poly-lysine-coated coverslips in EMMG medium.

### Fluorescence *in situ* hybridisation

Cells were grown to an OD_600_ of 0.5-0.7. For heat shock samples, 5.5 ml cell culture were mixed with 3.3 m pre-warmed medium, then shifted to 42°C for 1 hour under agitation. Cells were fixed by the addition of 1.25 ml 37% formaldehyde at 42°C for 20 min, then for an additional 1.5 h at RT on a roller mixer. For non-heat shock samples, 8.75 ml cell culture were fixed with 1.25 ml 37% formaldehyde for 1.5 h at RT. Cells were harvested (5 min, 3000 rpm) and washed with 5 ml 0.1 M KPO_4_, pH 6.4. Cells were then transferred to 1.5 ml tubes and incubated in 1 ml wash buffer (0.1 M KPO_4_ with 1.2 M sorbitol) at 4°C overnight. Cells were spheroplasted in 200 μl wash buffer containing 100 μg zymolyase T100 (Carl Roth) for 1 hour at 37°C, then washed once with 1 ml wash buffer. Cells were incubated in 200 µl 2 x SSC at RT for 10 minutes, then resuspended in pre-hybridisation buffer (50% formamide, 10% dextran sulphate, 125 μg/ml *E. coli* tRNA, 500 μg/ml herring sperm DNA, 1 x Denhardt’s solution) and incubated for 1 hour at 37°C. 1 µl of 1 pmol/µl Cy3-labelled oligo-d(T)_50_ probe was added to the sample mix and incubated at 37°C overnight in the dark. Cells were pelleted, gently resuspended in 600 µl of 0.5 x SSC and incubated on a rotating wheel for 30 minutes in the dark at RT. Cells were washed with 600 µl PBS, then resuspended in 50 µl PBS containing 0.1% NaN_3_. ½ volume was mounted on a poly-lysine-coated coverslip and left to sit for 30 min at RT in the dark. Non-adherent cells were removed by aspiration and cells then imaged under ROTIMount FluorCare DAPI (Carl Roth) using a Deltavision Ultra High-Resolution Microscope (Cytiva).

Image quantification was carried out with Fiji using a semiautomated ImageJ macro script (Suppl. File 1). In short, cellular segmentation was carried out by thresholding on the transmitted light channel (cell outlines) and DAPI stain (nuclei). Measurements of oligo-dT_50_ signal were performed on average intensity Z-projections for nucleus and cytoplasm (= total cell without nucleus) and the ratio of mean nuclear fluorescence intensity over mean cytoplasmic fluorescence intensity calculated for each cell. The displayed p-values for the pair-wise comparisons were calculated using the Wilcoxon test and the ggpubr package in R (Kassambara, 2023).

### SDS lysis and extract preparation

100 ml cultures were grown to an OD_600_ of 1.0 and harvested by centrifugation (2 min, 3.000xg at 4°C). Cell were washed with 1 ml ice-cold H_2_O and transferred into 2 ml microcentrifuge tubes before lysis with 400 µl glass beads (425-600 µm Ø) in 150 µl lysis buffer (10 mM Tris-HCl pH 8.0, 200 mM KCl, 2.5 mM MgCl_2_, 0.5 mM EDTA pH 8.0, 0.5% NP-40, protease inhibitor cocktail (Sigma Merck, P8215), 1mM PMSF) by vortexing for 20 min with 30 s on/off time intervals on ice. Lysates were transferred to a fresh microcentrifuge tube and cleared by centrifugation (5 min, 21,000xg at 4 °C). Protein concentration was determined by BCA assay (Novagen BCA Protein Assay Kit) on a NanoDrop Microvolume Spectrophotometer (Fisher Scientific). 10 µg total protein per lane were resolved on a 10% SDS-PAGE gel.

The following antibodies were used to detect proteins after membrane transfer: Anti-c-Myc, clone 9E10 (biomol, NSJ-F52056), anti-GFP from mouse IgG1κ (clones 7.1 and 13.1) (Sigma, 11814460001), anti-mini-AID-tag mAb, IgG2aκ, mouse monoclonal (Biozol, MBL-M214-3), anti-GAPDH, clone GA1R (biomol, MM-0163-P).

### Co-immunoprecipitation assays

Lysates were prepared from 240 OD_600_ units as described above. 1 mg total proteins were incubated with 15 µl pre-equilibrated magnetic GFP-TRAP beads (Chromotek) for 2 h on a rotator at 4 °C. Beads were washed 6 x with 1 ml lysis buffer for 5 min at 4 °C. Bound protein was then eluted with 60 µl HU loading buffer (8 M urea, 5% SDS, 200 mM Tris-HCl pH 6.5, 20 mM dithiothreitol (DTT), 1.5 mM bromophenol blue) for 5 min at 95°C. 20 µl per co-IP and 10 µl of input (1:10) were separated on a 10% SDS-PAGE gel.

### TCA lysis and extract preparation

Trichloroacetic acid (TCA) lysis was essentially performed as described (Mehta et al., 2010). 50 ml cultures were grown to an OD_600_ of 0.5-0.6OD_600_/mL and harvested by centrifugation (2 min, 3.000xg at 4°C). Cells were washed with 1 ml ice-cold H_2_O and transferred into 2 ml microcentrifuge tubes. Pellets were washed once with ice cold 20% TCA (15 s, 21,000xg at 4°C), and then lysed with 500 µl glass beads (425-600 µm Ø) in 500µl 20% TCA by vortexing 3 x 1 min with 1 min breaks on ice. The bottom of the tube was punctured with a hot 22-gauge ½ inch needle and lysates were transferred into new 1.5 ml microcentrifuge tubes by spinning (2min, 376xg at 4°C). Beads were washed once with 600 µl ice-cold 5% TCA and the wash pooled with the lysates. Precipitated proteins were pelleted by centrifugation (10 min, 18,000xg at 4°C). Pellets were washed with 1 ml ice-cold 100% EtOH (5 min, 21,000xg at 4°C). Pellets were resuspended in 84 µl 1M Tris pH 10.0 and 167 µl HU loading buffer (8M Urea, 5% SDS, 200 mM Tris-HCl pH 6.5, 20 mM DTT, 1.5 mM bromophenol blue) and boiled for 5 minutes at 95°C before samples were separation on a 10% SDS-PAGE gel (15µl per lane).

### RNA preparation and Northern Blot

RNA was prepared with the hot phenol method and resolved on 1.2% agarose gels after glyoxylation as described (Cruz & Houseley, 2020). For strand-specific Northern blot, digoxigenin (DIG)-labelled probes were in vitro-transcribed with MAXI-script (Ambion) and detected using the DIG system (Roche). Oligos for probe preparation are listed in Table S3. In all cases, methylene blue staining of ribosomal bands served as a loading control.

### RNA sequencing

For RNA sequencing, pellets from *S. pombe* and *S. cerevisiae* cultures were pooled in a 5:1 OD_600_ ratio immediately before cell lysis. Total RNA was extracted with the hot phenol method. Libraries were prepared with the VAHTS Stranded mRNA-seq Library Prep Kit (Vazyme Biotech) after poly(A) selection and sequenced on a NextSeq 2000 system (Illumina) or using the TruSeq protocol (Illumina) after ribodepletion with the Ribo-Zero Magnetic Kit for yeast (Epicentre) and sequenced on an Illumina HiSeq 4000 instrument.

### High-throughput sequencing data analysis

After read trimming with Trimmomatic, RNA-seq data were aligned to versions ASM294v2 and R64-1-1 of the *S. pomb*e and *S. cerevisiae* genome using the STAR aligner on the Galaxy server (Afgan et al., 2022; Bolger et al., 2014; Dobin et al., 2013). All reads aligning to both yeasts were filtered out using SAMtools, and the total number of reads aligning to *S. cerevisiae* used for normalization of the *S. pombe* data (H. Li et al., 2009). Differential gene expression analysis was carried out with the DESeq2 tool after read counting with HTSeq-count in Union mode on the ASM294v2.57 gene model provided by EnsemblFungi (Anders et al., 2015; Love et al., 2014; Martin et al., 2023). ChIP-seq data were aligned with BowTie2 (Langmead et al., 2009) and processed as above. Genomic ranges for metagene plots and heatmaps were selected using the packages rtracklayer and GenomicAlignments in R, and metagenes generated with the metagene2 package without normalization before geometric scaling (Fournier et al., 2022; Lawrence et al., 2009, 2013). GO term annotations were retrieved using biomaRt (Durinck et al., 2009). Plots were generated with the ggplot2 package in R (Wickham, 2016). For box plots, the lower and upper hinges correspond to the first and third quartiles, and the whiskers extend from the hinge to the smallest and largest value no further than 1.5 * IQR from the hinge (where IQR is the inter-quartile range). Outliers are plotted individually.

### Interaction prediction

Interaction between Srp2 and Rpb8 was predicted with Colabfold (Mirdita et al., 2022), which couples AlphaFold2 with Google Collaboratory, using model type ‘Alphafold2-multimer-v2’ with default parameters.

### Data accessibility

Raw (fastq) and processed (bedgraph) sequencing data can be downloaded from EBI ArrayExpress with the accession numbers E-MTAB-13712, E-MTAB-13714 and E-MTAB-13717. The mass spectrometry proteomics data have been deposited to the ProteomeXchange Consortium via the PRIDE (Deutsch et al., 2023) partner repository with the dataset identifier PXD048560.

## Supporting information

Supplemental Tables 2-4: strains, oligos, plasmids

Supplemental File: FISH quant ImageJ script

## Acknowledgments

We thank the National BioResource Project Japan, François Bachand, Tomoyasu Sugiyama, and Lidia Vasiljeva (LV) for strains and constructs. This work was supported by the Emmy Noether Programme and the priority programme SPP1935 of the DFG (KI1657/2-1 and KI1657/3-1) to CK. We are grateful to LV for support and advice, to Johanna Seidler for input on the FISH quantification script, and to Andreas Diepold, Vera Bettenworth and members of the lab for critical comments on the manuscript.

## References

Afgan E, Nekrutenko A, Grüning BA, Blankenberg D, Goecks J, Schatz MC, Ostrovsky AE, Mahmoud A, Lonie AJ, Syme A, Fouilloux A, Bretaudeau A, Nekrutenko A, Kumar A, Eschenlauer AC, Desanto AD, Guerler A, Serrano-Solano B, Batut B, Grüning BA, Langhorst BW, Carr B, Raubenolt BA, Hyde CJ, Bromhead CJ, Barnett CB, Royaux C, Gallardo C, Blankenberg D, Fornika DJ, Baker D, Bouvier D, Clements D, De Lima Morais DA, Tabernero DL, Lariviere D, Nasr E, Afgan E, Zambelli F, Heyl F, Psomopoulos F, Coppens F, Price GR, Cuccuru G, Corguillé G Le, Von Kuster G, Akbulut GG, Rasche H, Hans-Rudolf H, Eguinoa I, Makunin I, Ranawaka IJ, Taylor JP, Joshi J, Hillman-Jackson J, Chilton JM, Kamali K, Suderman K, Poterlowicz K, Yvan LB, Lopez-Delisle L, Sargent L, Bassetti ME, Tangaro MA, Van Den Beek M, Cech M, Bernt M, Fahrner M, Tekman M, Föll MC, Schatz MC, Crusoe MR, Roncoroni M, Kucher N, Coraor N, Stoler N, Rhodes N, Soranzo N, Pinter N, Goonasekera NA, Moreno PA, Videm P, Melanie P, Mandreoli P, Jagtap PD, Gu Q, Weber RJM, Lazarus R, Vorderman RHP, Hiltemann S, Golitsynskiy S, Garg S, Bray SA, Gladman SL, Leo S, Mehta SP, Griffin TJ, Jalili V, Yves V, Wen V, Nagampalli VK, Bacon WA, De Koning W, Maier W & Briggs PJ (2022) The Galaxy platform for accessible, reproducible and collaborative biomedical analyses: 2022 update. Nucleic Acids Res 50, W345–W351.

Ahn S-H, Kim M & Buratowski S (2004) Phosphorylation of serine 2 within the RNA polymerase II C-terminal domain couples transcription and 3’ end processing. Mol. Cell 13, 67–76.

Ahrné E, Molzahn L, Glatter T & Schmidt A (2013) Critical assessment of proteome-wide label-free absolute abundance estimation strategies. Proteomics 13, 2567–2578.

Albulescu LO, Sabet N, Gudipati M, Stepankiw N, Bergman ZJ, Huffaker TC & Pleiss JA (2012) A Quantitative, High-Throughput Reverse Genetic Screen Reveals Novel Connections between Pre–mRNA Splicing and 5′ and 3′ End Transcript Determinants. PLoS Genet 8, e1002530.

Anders S, Pyl PT & Huber W (2015) HTSeq—a Python framework to work with high-throughput sequencing data. Bioinformatics 31, 166–169.

Andersen CBF, Ballut L, Johansen JS, Chamieh H, Nielsen KH, Oliveira CLP, Pedersen JS, Séraphin B, Hir H Le & Andersen GR (2006) Structure of the exon junction core complex with a trapped DEAD-Box ATPase bound to RNA. Science (1979) 313, 1968–1972.

Assenholt J, Mouaikel J, Andersen KR, Brodersen DE, Libri D & Jensen TH (2008) Exonucleolysis is required for nuclear mRNA quality control in yeast THO mutants. RNA 14, 2305–13.

Atkinson SR, Marguerat S, Bitton DA, Rodríguez-López M, Rallis C, Lemay JF, Cotobal C, Malecki M, Smialowski P, Mata J, Korber P, Bachand F & Bähler J (2018) Long noncoding RNA repertoire and targeting by nuclear exosome, cytoplasmic exonuclease, and RNAi in fission yeast. RNA 24, 1195–1213.

Bähler J, Wu JQ, Longtine MS, Shah NG, McKenzie A, Steever AB, Wach A, Philippsen P & Pringle JR (1998) Heterologous modules for efficient and versatile PCR-based gene targeting in Schizosaccharomyces pombe. Yeast 14, 943–51.

Baltz AG, Munschauer M, Schwanhäusser B, Vasile A, Murakawa Y, Schueler M, Youngs N, Penfold-Brown D, Drew K, Milek M, Wyler E, Bonneau R, Selbach M, Dieterich C & Landthaler M (2012) The mRNA-bound proteome and its global occupancy profile on protein-coding transcripts. Mol Cell 46, 674–90.

Bao X, Guo X, Yin M, Tariq M, Lai Y, Kanwal S, Zhou J, Li N, Lv Y, Pulido-Quetglas C, Wang X, Ji L, Khan MJ, Zhu X, Luo Z, Shao C, Lim D-H, Liu X, Li N, Wang W, He M, Liu Y-L, Ward C, Wang T, Zhang G, Wang D, Yang J, Chen Y, Zhang C, Jauch R, Yang Y-G, Wang Y, Qin B, Anko M-L, Hutchins AP, Sun H, Wang H, Fu X-D, Zhang B & Esteban MA (2018) Capturing the interactome of newly transcribed RNA. Nat Methods 15, 213–220.

Barta I & Iggo RD (1995) Autoregulation of expression of the yeast Dbp2p “DEAD-box” protein is mediated by sequences in the conserved DBP2 intron. EMBO J 14, 3800–8.

Bazan JF (2008) An old HAT in human p300/CBP and yeast Rtt109. Cell Cycle 7, 1884–1886.

Beckmann BM, Horos R, Fischer B, Castello A, Eichelbaum K, Alleaume A-M, Schwarzl T, Curk T, Foehr S, Huber W, Krijgsveld J & Hentze MW (2015) The RNA-binding proteomes from yeast to man harbour conserved enigmRBPs. Nat Commun 6, 10127.

Birot A, Kus K, Priest E, Al Alwash A, Castello A, Mohammed S, Vasiljeva L & Kilchert C (2021) RNA-binding protein Mub1 and the nuclear RNA exosome act to fine-tune environmental stress response. Life Sci Alliance 5, e202101111.

Bolger AM, Lohse M & Usadel B (2014) Trimmomatic: a flexible trimmer for Illumina sequence data. Bioinformatics 30, 2114–2120.

Boreikaitė V & Passmore LA (2023) 3’-End Processing of Eukaryotic mRNA: Machinery, Regulation, and Impact on Gene Expression. Annu Rev Biochem 92.

Bresson SM, Hunter O V., Hunter AC & Conrad NK (2015) Canonical Poly(A) Polymerase Activity Promotes the Decay of a Wide Variety of Mammalian Nuclear RNAs T. Jensen, ed. PLoS Genet 11, e1005610.

Buszczak M & Spradling AC (2006) The Drosophila P68 RNA helicase regulates transcriptional deactivation by promoting RNA release from chromatin. Genes Dev 20, 977–989.

Del Campo M & Lambowitz AM (2009) Structure of the Yeast DEAD Box Protein Mss116p Reveals Two Wedges that Crimp RNA. Mol Cell 35, 598–609.

Carminati M, Rodríguez-Molina JB, Manav MC, Bellini D & Passmore LA (2023) A direct interaction between CPF and RNA Pol II links RNA 3′ end processing to transcription. Mol Cell 83, 4461–4478.e13.

Castello A, Fischer B, Eichelbaum K, Horos R, Beckmann BM, Strein C, Davey NE, Humphreys DT, Preiss T, Steinmetz LM, Krijgsveld J & Hentze MW (2012) Insights into RNA biology from an atlas of mammalian mRNA-binding proteins. Cell 149, 1393–406.

Chen H-M, Futcher B & Leatherwood J (2011) The fission yeast RNA binding protein Mmi1 regulates meiotic genes by controlling intron specific splicing and polyadenylation coupled RNA turnover. PLoS One 6, e26804.

Cheng H, Dufu K, Lee C-S, Hsu JL, Dias A & Reed R (2006) Human mRNA Export Machinery Recruited to the 5′ End of mRNA. Cell 127, 1389–1400.

Cloutier SC, Wang S, Ma WK, Al Husini N, Dhoondia Z, Ansari A, Pascuzzi PE & Tran EJ (2016) Regulated Formation of lncRNA-DNA Hybrids Enables Faster Transcriptional Induction and Environmental Adaptation. Mol Cell 61, 393–404.

Cortazar MA, Sheridan RM, Erickson B, Fong N, Glover-Cutter K, Brannan K & Bentley DL (2019) Control of RNA Pol II Speed by PNUTS-PP1 and Spt5 Dephosphorylation Facilitates Termination by a “Sitting Duck Torpedo” Mechanism. Mol Cell 76, 896–908.e4.

Cox J & Mann M (2008) MaxQuant enables high peptide identification rates, individualized p.p.b.-range mass accuracies and proteome-wide protein quantification. Nature Biotechnology 2008 26:12 26, 1367–1372.

Cristini A, Groh M, Kristiansen MS & Gromak N (2018) RNA/DNA Hybrid Interactome Identifies DXH9 as a Molecular Player in Transcriptional Termination and R-Loop-Associated DNA Damage. Cell Rep 23, 1891–1905.

Cruz C & Houseley J (2020) Protocols for Northern Analysis of Exosome Substrates and Other Noncoding RNAs. Methods Mol Biol 2062, 83–103.

Custódio N, Carmo-Fonseca M, Geraghty F, Pereira HS, Grosveld F & Antoniou M (1999) Inefficient processing impairs release of RNA from the site of transcription. EMBO J 18, 2855–66.

Custódio N, Vivo M, Antoniou M & Carmo-Fonseca M (2007) Splicing- and cleavage-independent requirement of RNA polymerase II CTD for mRNA release from the transcription site. J Cell Biol 179, 199–207.

Dardenne E, Polay Espinoza M, Fattet L, Germann S, Lambert M-P, Neil H, Zonta E, Mortada H, Gratadou L, Deygas M, Chakrama FZ, Samaan S, Desmet F-O, Tranchevent L-C, Dutertre M, Rimokh R, Bourgeois CF & Auboeuf D (2014) RNA helicases DDX5 and DDX17 dynamically orchestrate transcription, miRNA, and splicing programs in cell differentiation. Cell Rep 7, 1900–13.

Deutsch EW, Bandeira N, Perez-Riverol Y, Sharma V, Carver JJ, Mendoza L, Kundu DJ, Wang S, Bandla C, Kamatchinathan S, Hewapathirana S, Pullman BS, Wertz J, Sun Z, Kawano S, Okuda S, Watanabe Y, Maclean B, Maccoss MJ, Zhu Y, Ishihama Y & Vizcaíno JA (2023) The ProteomeXchange consortium at 10 years: 2023 update. Nucleic Acids Res 51, D1539–D1548.

Ding D-Q, Okamasa K, Yoshimura Y, Matsuda A, Yamamoto TG, Hiraoka Y, Nakayama J & Da-Qiao Ding C (2023) The mechanism of homologous chromosome recognition and pairing facilitated by chromosome-tethered protein-RNA condensates. bioRxiv 2023.12.24.573283.

Dobin A, Davis CA, Schlesinger F, Drenkow J, Zaleski C, Jha S, Batut P, Chaisson M & Gingeras TR (2013) STAR: ultrafast universal RNA-seq aligner. Bioinformatics 29, 15–21.

Donsbach P & Klostermeier D (2021) Regulation of RNA helicase activity: Principles and examples. Biol Chem 402, 529–559.

Durinck S, Spellman PT, Birney E & Huber W (2009) Mapping identifiers for the integration of genomic datasets with the R/Bioconductor package biomaRt. Nat Protoc 4, 1184–91.

Eckmann CR, Rammelt C & Wahle E (2011) Control of poly(A) tail length. Wiley Interdiscip Rev RNA 2, 348–361.

Egan ED, Braun CR, Gygi SP & Moazed D (2014) Post-transcriptional regulation of meiotic genes by a nuclear RNA silencing complex. RNA 20, 867–81.

Eick D & Geyer M (2013) The RNA polymerase II carboxy-terminal domain (CTD) code. Chem Rev 113, 8456–8490.

Enders M, Neumann P, Dickmanns A & Ficner R (2023) Structure and function of spliceosomal DEAH-box ATPases. Biol Chem 404, 851–866.

Fairman ME, Maroney PA, Wang W, Bowers HA, Gollnick P, Nilsen TW & Jankowsy E (2004) Protein Displacement by DExH/D “RNA Helicases” Without Duplex Unwinding. Science (1979) 304, 730–734.

Fan J, Kuai B, Wang K, Wang L, Wang Y, Wu X, Chi B, Li G & Cheng H (2018) mRNAs are sorted for export or degradation before passing through nuclear speckles. Nucleic Acids Res 46, 8404–8416.

Fong N, Brannan K, Erickson B, Kim H, Cortazar MA, Sheridan RM, Nguyen T, Karp S & Bentley DL (2015) Effects of Transcription Elongation Rate and Xrn2 Exonuclease Activity on RNA Polymerase II Termination Suggest Widespread Kinetic Competition. Mol Cell 60, 256–267.

Fong N, Öhman M & Bentley DL (2009) Fast ribozyme cleavage releases transcripts from RNA polymerase II and aborts co-transcriptional pre-mRNA processing. Nat. Struct. Mol. Biol. 16, 916–22.

Fournier E, Joly Beauparlant C, Lippens C & Droit A (2022) metagene2: A package to produce metagene plots. R package version 1.14.0.

Gallardo P, Real-Calderón P, Flor-Parra I, Salas-Pino S & Daga RR (2020) Acute Heat Stress Leads to Reversible Aggregation of Nuclear Proteins into Nucleolar Rings in Fission Yeast. Cell Rep 33.

Glatter T, Ludwig C, Ahrné E, Aebersold R, Heck AJR & Schmidt A (2012) Large-scale quantitative assessment of different in-solution protein digestion protocols reveals superior cleavage efficiency of tandem Lys-C/trypsin proteolysis over trypsin digestion. J Proteome Res 11, 5145–5156.

Gruber AJ & Zavolan M (2019) Alternative cleavage and polyadenylation in health and disease. Nat Rev Genet 20, 599–614.

Hagan IM (2016) Immunofluorescence Microscopy of Schizosaccharomyces pombe Using Chemical Fixation. Cold Spring Harb Protoc 2016, pdb.prot091017.

Hammell CM, Gross S, Zenklusen D, Heath C V, Stutz F, Moore C & Cole CN (2002) Coupling of termination, 3’ processing, and mRNA export. Mol Cell Biol 22, 6441–57.

Harigaya Y, Tanaka H, Yamanaka S, Tanaka K, Watanabe Y, Tsutsumi C, Chikashige Y, Hiraoka Y, Yamashita A & Yamamoto M (2006) Selective elimination of messenger RNA prevents an incidence of untimely meiosis. Nature 442, 45–50.

Harris MA, Rutherford KM, Hayles J, Lock A, Bähler J, Oliver SG, Mata J & Wood V (2022) Fission stories: using PomBase to understand Schizosaccharomyces pombe biology. Genetics 220.

Henfrey C, Murphy S & Tellier M (2023) Regulation of mature mRNA levels by RNA processing efficiency. NAR Genom Bioinform 5.

Henn A, Bradley MJ & De La Cruz EM (2012) ATP Utilization and RNA Conformational Rearrangement by DEAD-Box Proteins. Annual Review of Biophysics 41, 247–267.

Herrera-Moyano E, Mergui X, García-Rubio ML, Barroso S & Aguilera A (2014) The yeast and human FACT chromatin-reorganizing complexes solve R-loop-mediated transcription–replication conflicts. Genes Dev 28, 735–748.

Herzel L, Straube K & Neugebauer KM (2018) Long-read sequencing of nascent RNA reveals coupling among RNA processing events. Genome Res 28, 1008–1019.

Hilbert M, Karow AR & Klostermeier D (2009) The mechanism of ATP-dependent RNA unwinding by DEAD box proteins. Biol Chem 390, 1237–1250.

Hilleren P, McCarthy T, Rosbash M, Parker R & Jensen TH (2001) Quality control of mRNA 3’-end processing is linked to the nuclear exosome. Nature 413, 538–42.

Hirose T, Ninomiya K, Nakagawa S & Yamazaki T (2022) A guide to membraneless organelles and their various roles in gene regulation. Nature Reviews Molecular Cell Biology 2022 24:4 24, 288–304.

Hondele M, Sachdev R, Heinrich S, Wang J, Vallotton P, Fontoura BMA & Weis K (2019) DEAD-box ATPases are global regulators of phase-separated organelles. Nature 573, 144–148.

Jimeno S, Rondón AG, Luna R & Aguilera A (2002) The yeast THO complex and mRNA export factors link RNA metabolism with transcription and genome instability. EMBO J 21, 3526–3535.

Kaida D, Berg MG, Younis I, Kasim M, Singh LN, Wan L & Dreyfuss G (2010) U1 snRNP protects pre-mRNAs from premature cleavage and polyadenylation. Nature 468, 664–8.

Kammers K, Cole RN, Tiengwe C & Ruczinski I (2015) Detecting significant changes in protein abundance. EuPA Open Proteom 7, 11–19.

Kar A, Fushimi K, Zhou X, Ray P, Shi C, Chen X, Liu Z, Chen S & Wu JY (2011) RNA Helicase p68 (DDX5) Regulates tau Exon 10 Splicing by Modulating a Stem-Loop Structure at the 5′ Splice Site. Mol Cell Biol 31, 1812–1821.

Kassambara A (2023) ggpubr: ‘ggplot2’ Based Publication Ready Plots. – Available at: https://rdrr.io/cran/ggpubr/

Katahira J, Senokuchi K & Hieda M (2020) Human THO maintains the stability of repetitive DNA. Genes to Cells pgtc.12760.

Kecman T, Heo DH & Vasiljeva L (2018) Profiling RNA Polymerase II Phosphorylation Genome-Wide in Fission Yeast. Methods Enzymol 612, 489–504.

Kecman T, Kuś K, Heo D-H, Duckett K, Birot A, Liberatori S, Mohammed S, Geis-Asteggiante L, Robinson C V. & Vasiljeva L (2018) Elongation/Termination Factor Exchange Mediated by PP1 Phosphatase Orchestrates Transcription Termination. Cell Rep 25, 259–269.e5.

Kilchert C (2020) RNA Exosomes and Their Cofactors. Methods Mol Biol 2062, 215–235.

Kilchert C, Kecman T, Priest E, Hester S, Aydin E, Kus K, Rossbach O, Castello A, Mohammed S & Vasiljeva L (2020) System-wide analyses of the fission yeast poly(A)+RNA interactome reveal insights into organization and function of RNA-protein complexes. Genome Res 30, 1012–1026.

Kilchert C, Kecman T, Priest E, Hester S, Kus K, Castello A, Mohammed S & Vasiljeva L (2019) System-wide analyses of the fission yeast poly(A)+ RNA interactome reveal insights into organisation and function of RNA-protein complexes. bioRxiv 748194.

Kilchert C, Wittmann S, Passoni M, Shah S, Granneman S & Vasiljeva L (2015) Regulation of mRNA levels by decay-promoting introns that recruit the exosome specificity factor Mmi1. Cell Rep 13, 2504–2515.

Kilchert C, Wittmann S & Vasiljeva L (2016) The regulation and functions of the nuclear RNA exosome complex. Nat Rev Mol Cell Biol 17, 227–239.

Kim HD, Choe J & Seo YS (1999) The sen1(+) gene of Schizosaccharomyces pombe, a homologue of budding yeast SEN1, encodes an RNA and DNA helicase. Biochemistry 38, 14697–14710.

Kim M, Krogan NJ, Vasiljeva L, Rando OJ, Nedea E, Greenblatt JF & Buratowski S (2004) The yeast Rat1 exonuclease promotes transcription termination by RNA polymerase II. Nature 432, 517–22.

Kim M, Vasiljeva L, Rando OJ, Zhelkovsky A, Moore C & Buratowski S (2006) Distinct pathways for snoRNA and mRNA termination. Mol. Cell 24, 723–34.

Knop M, Siegers K, Pereira G, Zachariae W, Winsor B, Nasmyth K & Schiebel E (1999) Epitope Tagging of Yeast Genes using a PCR-based Strategy: More Tags and Improved Practical Routines. Yeast 15, 963–972.

Kyburz A, Friedlein A, Langen H & Keller W (2006) Direct Interactions between Subunits of CPSF and the U2 snRNP Contribute to the Coupling of Pre-mRNA 3′ End Processing and Splicing. Mol Cell 23, 195–205.

Lai YH, Choudhary K, Cloutier SC, Xing Z, Aviran S & Tran EJ (2019) Genome-Wide Discovery of DEAD-Box RNA Helicase Targets Reveals RNA Structural Remodeling in Transcription Termination. Genetics 212, 153–174.

Langmead B, Trapnell C, Pop M & Salzberg SL (2009) Ultrafast and memory-efficient alignment of short DNA sequences to the human genome. Genome Biol 10, 1–10.

Larochelle M, Robert MA, Hébert JN, Liu X, Matteau D, Rodrigue S, Tian B, Jacques PÉ & Bachand F (2018) Common mechanism of transcription termination at coding and noncoding RNA genes in fission yeast. Nature Communications 2018 9:1 9, 1–15.

Lawrence M, Gentleman R & Carey V (2009) rtracklayer: an R package for interfacing with genome browsers. Bioinformatics 25, 1841–1842.

Lawrence M, Huber W, Pagès H, Aboyoun P, Carlson M, Gentleman R, Morgan MT & Carey VJ (2013) Software for computing and annotating genomic ranges. PLoS Comput Biol 9.

Ledoux S & Guthrie C (2011) Regulation of the Dbp5 ATPase cycle in mRNP remodeling at the nuclear pore: a lively new paradigm for DEAD-box proteins. Genes Dev 25, 1109–14.

Lee ES, Smith HW, Wolf EJ, Guvenek A, Wang YE, Emili A, Tian B & Palazzo AF (2022) ZFC3H1 and U1-70K promote the nuclear retention of mRNAs with 5’ splice site motifs within nuclear speckles. RNA rna.079104.122.

Lee NN, Chalamcharla VR, Reyes-Turcu FE, Mehta S, Zofall M, Balachandran V, Dhakshnamoorthy J, Taneja N, Yamanaka S, Zhou M & Grewal SIS (2013) Mtr4-like protein coordinates nuclear RNA processing for heterochromatin assembly and for telomere maintenance. Cell 155, 1061–74.

Lemay J-F, Larochelle M, Marguerat S, Atkinson S, Bähler J & Bachand F (2014) The RNA exosome promotes transcription termination of backtracked RNA polymerase II. Nat Struct Mol Biol 21, 919–926.

Lemay J-F, Marguerat S, Larochelle M, Liu X, van Nues R, Hunyadkürti J, Hoque M, Tian B, Granneman S, Bähler J & Bachand F (2016) The Nrd1-like protein Seb1 coordinates cotranscriptional 3′ end processing and polyadenylation site selection. Genes Dev 30, 1558–1572.

Li H, Handsaker B, Wysoker A, Fennell T, Ruan J, Homer N, Marth G, Abecasis G & Durbin R (2009) The Sequence Alignment/Map format and SAMtools. Bioinformatics 25, 2078–2079.

Li J, Querl L, Coban I, Salinas G & Krebber H (2023) Surveillance of 3′ mRNA cleavage during transcription termination requires CF IB/Hrp1. Nucleic Acids Res 51, 8758–8773.

Li L, Roy K, Katyal S, Sun X, Bléoo S & Godbout R (2006) Dynamic nature of cleavage bodies and their spatial relationship to DDX1 bodies, Cajal bodies, and gems. Mol Biol Cell 17, 1126–1140.

Lin C, Yang L, Yang JJ, Huang Y & Liu Z-R (2005) ATPase/helicase activities of p68 RNA helicase are required for pre-mRNA splicing but not for assembly of the spliceosome. Mol Cell Biol 25, 7484–93.

Linder P (2006) Dead-box proteins: a family affair—active and passive players in RNP-remodeling. Nucleic Acids Res 34, 4168–4180.

Linder P & Lasko P (2006) Bent out of Shape: RNA Unwinding by the DEAD-Box Helicase Vasa. Cell 125, 219–221.

Liu Y, Warfield L, Zhang C, Luo J, Allen J, Lang WH, Ranish J, Shokat KM & Hahn S (2009) Phosphorylation of the transcription elongation factor Spt5 by yeast Bur1 kinase stimulates recruitment of the PAF complex. Mol Cell Biol 29, 4852–4863.

Love MI, Huber W & Anders S (2014) Moderated estimation of fold change and dispersion for RNA-seq data with DESeq2. Genome Biol 15, 550.

Lund MK & Guthrie C (2005) The DEAD-Box Protein Dbp5p Is Required to Dissociate Mex67p from Exported mRNPs at the Nuclear Rim. Mol Cell 20, 645–651.

Ma WK, Cloutier SC & Tran EJ (2013) The DEAD-box Protein Dbp2 Functions with the RNA-Binding Protein Yra1 to Promote mRNP Assembly. J Mol Biol 425, 3824–3838.

Ma WK, Paudel BP, Xing Z, Sabath IG, Rueda D & Tran EJ (2016) Recruitment, Duplex Unwinding and Protein-Mediated Inhibition of the Dead-Box RNA Helicase Dbp2 at Actively Transcribed Chromatin. J Mol Biol 428, 1091–1106.

Marayati BF, Hoskins V, Boger RW, Tucker JF, Fishman ES, Bray AS & Zhang K (2016) The fission yeast MTREC and EJC orthologs ensure the maturation of meiotic transcripts during meiosis. RNA 22, 1349–1359.

Martín Caballero L, Capella M, Barrales RR, Dobrev N, van Emden T, Hirano Y, Suma Sreechakram VN, Fischer-Burkart S, Kinugasa Y, Nevers A, Rougemaille M, Sinning I, Fischer T, Hiraoka Y & Braun S (2022) The inner nuclear membrane protein Lem2 coordinates RNA degradation at the nuclear periphery. Nature Structural & Molecular Biology 2022 29:9 29, 910–921.

Martin FJ, Amode MR, Aneja A, Austine-Orimoloye O, Azov AG, Barnes I, Becker A, Bennett R, Berry A, Bhai J, Bhurji SK, Bignell A, Boddu S, Branco Lins PR, Brooks L, Ramaraju SB, Charkhchi M, Cockburn A, Da Rin Fiorretto L, Davidson C, Dodiya K, Donaldson S, El Houdaigui B, El Naboulsi T, Fatima R, Giron CG, Genez T, Ghattaoraya GS, Martinez JG, Guijarro C, Hardy M, Hollis Z, Hourlier T, Hunt T, Kay M, Kaykala V, Le T, Lemos D, Marques-Coelho D, Marugán JC, Merino GA, Mirabueno LP, Mushtaq A, Hossain SN, Ogeh DN, Sakthivel MP, Parker A, Perry M, Piliota I, Prosovetskaia I, Perez-Silva JG, Salam AIA, Saraiva-Agostinho N, Schuilenburg H, Sheppard D, Sinha S, Sipos B, Stark W, Steed E, Sukumaran R, Sumathipala D, Suner MM, Surapaneni L, Sutinen K, Szpak M, Tricomi FF, Urbina-Gómez D, Veidenberg A, Walsh TA, Walts B, Wass E, Willhoft N, Allen J, Alvarez-Jarreta J, Chakiachvili M, Flint B, Giorgetti S, Haggerty L, Ilsley GR, Loveland JE, Moore B, Mudge JM, Tate J, Thybert D, Trevanion SJ, Winterbottom A, Frankish A, Hunt SE, Ruffier M, Cunningham F, Dyer S, Finn RD, Howe KL, Harrison PW, Yates AD & Flicek P (2023) Ensembl 2023. Nucleic Acids Res 51, D933–D941.

Masuda S, Das R, Cheng H, Hurt E, Dorman N & Reed R (2005) Recruitment of the human TREX complex to mRNA during splicing. Genes Dev 19, 1512–1517.

Matia-González AM, Laing EE & Gerber AP (2015) Conserved mRNA-binding proteomes in eukaryotic organisms. Nat Struct Mol Biol 22, 1027–33.

Matsuyama A, Arai R, Yashiroda Y, Shirai A, Kamata A, Sekido S, Kobayashi Y, Hashimoto A, Hamamoto M, Hiraoka Y, Horinouchi S & Yoshida M (2006) ORFeome cloning and global analysis of protein localization in the fission yeast Schizosaccharomyces pombe. Nat Biotechnol 24, 841–847.

Matsuyama A, Shirai A, Yashiroda Y, Kamata A, Horinouchi S & Yoshida M (2004) pDUAL, a multipurpose, multicopy vector capable of chromosomal integration in fission yeast. Yeast 21, 1289–305.

Mbogning J, Nagy S, Pagé V, Schwer B, Shuman S, Fisher RP & Tanny JC (2013) The PAF Complex and Prf1/Rtf1 Delineate Distinct Cdk9-Dependent Pathways Regulating Transcription Elongation in Fission Yeast. PLoS Genet 9, e1004029.

Mehta M, Braberg H, Wang S, Lozsa A, Shales M, Solache A, Krogan NJ & Keogh MC (2010) Individual Lysine Acetylations on the N Terminus of Saccharomyces cerevisiae H2A.Z Are Highly but Not Differentially Regulated. J Biol Chem 285, 39855.

Meola N, Domanski M, Karadoulama E, Chen Y, Gentil C, Pultz D, Vitting-Seerup K, Lykke-Andersen S, Andersen JS, Sandelin A & Jensen TH (2016) Identification of a Nuclear Exosome Decay Pathway for Processed Transcripts. Mol Cell 64, 520–533.

Mersaoui SY, Yu Z, Coulombe Y, Karam M, Busatto FF, Masson J-Y & Richard S (2019) Arginine methylation of the DDX5 helicase RGG/RG motif by PRMT5 regulates resolution of RNA:DNA hybrids. EMBO J 38, e100986.

Millevoi S, Loulergue C, Dettwiler S, Karaa SZ, Keller W, Antoniou M & Vagner S (2006) An interaction between U2AF 65 and CF Im links the splicing and 3′ end processing machineries. EMBO Journal 25, 4854–4864.

Mirdita M, Schütze K, Moriwaki Y, Heo L, Ovchinnikov S & Steinegger M (2022) ColabFold: making protein folding accessible to all. Nature Methods 2022 19:6 19, 679–682.

Mischo HE, Gómez-González B, Grzechnik P, Rondón AG, Wei W, Steinmetz LM, Aguilera A & Proudfoot NJ (2011) Yeast Sen1 helicase protects the genome from transcription-associated instability. Mol. Cell 41, 21–32.

Monahan BJ, Villén J, Marguerat S, Bähler J, Gygi SP & Winston F (2008) Fission yeast SWI/SNF and RSC complexes show compositional and functional differences from budding yeast. Nature Structural & Molecular Biology 2008 15:8 15, 873–880.

Montpetit B, Thomsen ND, Helmke KJ, Seeliger MA, Berger JM & Weis K (2011) A conserved mechanism of DEAD-box ATPase activation by nucleoporins and InsP6 in mRNA export. Nature 2011 472:7342 472, 238–242.

Mooney SM, Goel A, D’Assoro AB, Salisbury JL & Janknecht R (2010) Pleiotropic effects of p300-mediated acetylation on p68 and p72 RNA helicase. J Biol Chem 285, 30443–30452.

Moore MJ & Proudfoot NJ (2009) Pre-mRNA processing reaches back to transcription and ahead to translation. Cell 136, 688–700.

Moreno S, Klar A & Nurse P (1991) [56] Molecular genetic analysis of fission yeast Schizosaccharomyces pombe. In Methods in enzymology. pp.795–823.

Neve J, Patel R, Wang Z, Louey A & Furger AM (2017) Cleavage and polyadenylation: Ending the message expands gene regulation. RNA Biol 14, 865.

Niwa M, Rose SD & Berget SM (1990) In vitro polyadenylation is stimulated by the presence of an upstream intron. Genes Dev 4, 1552–1559.

Ozgur S, Buchwald G, Falk S, Chakrabarti S, Prabu JR & Conti E (2015) The conformational plasticity of eukaryotic RNA-dependent ATPases. FEBS Journal 282, 850–863.

Pacheco-Fiallos B, Vorländer MK, Riabov-Bassat D, Fin L, O’Reilly FJ, Ayala FI, Schellhaas U, Rappsilber J & Plaschka C (2023) mRNA recognition and packaging by the human transcription–export complex. Nature 2023 1–8.

Pandya-Jones A, Bhatt DM, Lin CH, Tong AJ, Smale ST & Black DL (2013) Splicing kinetics and transcript release from the chromatin compartment limit the rate of Lipid A-induced gene expression. RNA 19, 811.

Parua PK, Booth GT, Sansó M, Benjamin B, Tanny JC, Lis JT & Fisher RP (2018) A Cdk9-PP1 switch regulates the elongation-termination transition of RNA polymerase II. Nature 558, 460–464.

Penzo A, Dubarry M, Brocas C, Zheng M, Mangione RM, Rougemaille M, Goncalves C, Lautier O, Libri D, Simon MN, Géli V, Dubrana K & Palancade B (2023) A R-loop sensing pathway mediates the relocation of transcribed genes to nuclear pore complexes. Nature Communications 2023 14:1 14, 1–15.

Polenkowski M, Allister AB, Burbano de Lara S, Pierce A, Geary B, El Bounkari O, Wiehlmann L, Hoffmann A, Whetton AD, Tamura T & Tran DDH (2023) THOC5 complexes with DDX5, DDX17, and CDK12 to regulate R loop structures and transcription elongation rate. iScience 26, 105784.

Porrua O & Libri D (2015) Transcription termination and the control of the transcriptome: why, where and how to stop. Nat Rev Mol Cell Biol 16, 190–202.

Proudfoot NJ (2011) Ending the message: poly(A) signals then and now. Genes Dev 25, 1770–82.

Putnam AA & Jankowsky E (2013) DEAD-box helicases as integrators of RNA, nucleotide and protein binding. Biochimica et Biophysica Acta (BBA) - Gene Regulatory Mechanisms 1829, 884–893.

Qu X, Lykke-Andersen S, Nasser T, Saguez C, Bertrand E, Jensen TH & Moore C (2009) Assembly of an Export-Competent mRNP Is Needed for Efficient Release of the 3′-End Processing Complex after Polyadenylation. Mol Cell Biol 29, 5327–5338.

Rodríguez-Molina JB & Turtola M (2023) Birth of a poly(A) tail: mechanisms and control of mRNA polyadenylation. FEBS Open Bio 13, 1140–1153.

Rodríguez-Molina JB, West S & Passmore LA (2023) Knowing when to stop: Transcription termination on protein-coding genes by eukaryotic RNAPII. Mol Cell 83, 404–415.

Rossow KL & Janknecht R (2003) Synergism between p68 RNA helicase and the transcriptional coactivators CBP and p300. Oncogene 2003 22:1 22, 151–156.

Rougemaille M, Dieppois G, Kisseleva-Romanova E, Gudipati RK, Lemoine S, Blugeon C, Boulay J, Jensen TH, Stutz F, Devaux F & Libri D (2008) THO/Sub2p functions to coordinate 3’-end processing with gene-nuclear pore association. Cell 135, 308–21.

Saavedra C, Tuug KS, Amberg DC, Hopper AK & Cole CN (1996) Regulation of mRNA export in response to stress in Saccharomyces cerevisiae. Genes Dev 10, 1608–1620.

Saguez C, Schmid M, Olesen JR, Ghazy MAE-H, Qu X, Poulsen MB, Nasser T, Moore C & Jensen TH (2008) Nuclear mRNA surveillance in THO/sub2 mutants is triggered by inefficient polyadenylation. Mol. Cell 31, 91–103.

Sanso M & Fisher RP (2013) Modelling the CDK-dependent transcription cycle in fission yeast. Biochem Soc Trans 41, 1660–1665.

Schindelin J, Arganda-Carreras I, Frise E, Kaynig V, Longair M, Pietzsch T, Preibisch S, Rueden C, Saalfeld S, Schmid B, Tinevez JY, White DJ, Hartenstein V, Eliceiri K, Tomancak P & Cardona A (2012) Fiji: an open-source platform for biological-image analysis. Nature Methods 2012 9:7 9, 676–682.

Schul W, Groenhout B, Koberna K, Takagaki Y, Jenny A, Manders EM, Raska I, van Driel R & de Jong L (1996) The RNA 3’ cleavage factors CstF 64 kDa and CPSF 100 kDa are concentrated in nuclear domains closely associated with coiled bodies and newly synthesized RNA. EMBO J 15, 2883–92.

Schwer B (2001) A new twist on RNA helicases: DExH/D box proteins as RNPases. Nature Structural Biology 2001 8:2 8, 113–116.

Sengoku T, Nureki O, Nakamura A, Kobayashi S & Yokoyama S (2006) Structural Basis for RNA Unwinding by the DEAD-Box Protein Drosophila Vasa. Cell 125, 287–300.

Shi Y & Manley JL (2015) The end of the message: Multiple protein–RNA interactions define the mRNA polyadenylation site. Genes Dev 29, 889–897.

Shichino Y, Otsubo Y, Yamamoto M & Yamashita A (2020) Meiotic gene silencing complex MTREC/NURS recruits the nuclear exosome to YTH-RNA-binding protein Mmi1. PLoS Genet 16, e1008598.

Siam R, Dolan WP & Forsburg SL (2004) Choosing and using Schizosaccharomyces pombe plasmids. Methods 33, 189–198.

Silla T, Karadoulama E, Mąkosa D, Lubas M & Jensen TH (2018) The RNA Exosome Adaptor ZFC3H1 Functionally Competes with Nuclear Export Activity to Retain Target Transcripts. Cell Rep 23, 2199–2210.

Silla T, Schmid M, Dou Y, Garland W, Milek M, Imami K, Johnsen D, Polak P, Andersen JS, Selbach M, Landthaler M & Jensen TH (2020) The human ZC3H3 and RBM26/27 proteins are critical for PAXT-mediated nuclear RNA decay. Nucleic Acids Res 48, 2518–2530.

Smalec BM, Ietswaart R, Choquet K, McShane E, West ER & Churchman LS (2022) Genome-wide quantification of RNA flow across subcellular compartments reveals determinants of the mammalian transcript life cycle. bioRxiv 2022.08.21.504696.

Song X, Xu R & Sugiyama T (2021) Two plasmid modules for introducing the auxin-inducible degron into the fission yeast Schizosaccharomyces pombe by PCR-based gene targeting. MicroPubl Biol.

Soni K, Sivadas A, Horvath A, Dobrev N, Hayashi R, Kiss L, Simon B, Wild K, Sinning I & Fischer T (2023) Mechanistic insights into RNA surveillance by the canonical poly(A) polymerase Pla1 of the MTREC complex. Nature Communications 2023 14:1 14, 1–20.

Steinmetz EJ, Conrad NK, Brow D a & Corden JL (2001) RNA-binding protein Nrd1 directs poly(A)-independent 3’-end formation of RNA polymerase II transcripts. Nature 413, 327–31.

Sträßer K, Masuda S, Mason P, Pfannstiel J, Oppizzi M, Rodríguez-Navarro S, Rondón AG, Aguilera A, Struhl K, Reed R & Hurt E (2002) TREX is a conserved complex coupling transcription with messenger RNA export. Nature 417, 304–8.

Sugiyama T & Sugioka-Sugiyama R (2011) Red1 promotes the elimination of meiosis-specific mRNAs in vegetatively growing fission yeast. EMBO J 30, 1027–39.

Sugiyama T, Sugioka-Sugiyama R, Hada K & Niwa R (2012) Rhn1, a nuclear protein, is required for suppression of meiotic mRNAs in mitotically dividing fission yeast. PLoS One 7, e42962.

Sugiyama T, Wanatabe N, Kitahata E, Tani T & Sugioka-Sugiyama R (2013) Red5 and three nuclear pore components are essential for efficient suppression of specific mRNAs during vegetative growth of fission yeast. Nucleic Acids Res 41, 6674–86.

Tang Z, Käufer NF & Lin RJ (2002) Interactions between two fission yeast serine/arginine-rich proteins and their modulation by phosphorylation. Biochem J 368, 527–534.

Tashiro S, Asano T, Kanoh J & Ishikawa F (2013) Transcription-induced chromatin association of RNA surveillance factors mediates facultative heterochromatin formation in fission yeast. Genes Cells 18, 327–39.

Tedeschi FA, Cloutier SC, Tran EJ & Jankowsky E (2018) The DEAD-box protein Dbp2p is linked to noncoding RNAs, the helicase Sen1p, and R-loops. RNA 24, 1693–1705.

Terrone S, Valat J, Fontrodona N, Giraud G, Claude J-B, Combe E, Lapendry A, Polvèche H, Ameur L Ben, Duvermy A, Modolo L, Bernard P, Mortreux F, Auboeuf D & Bourgeois CF (2022) RNA helicase-dependent gene looping impacts messenger RNA processing. Nucleic Acids Res 50, 9226–9246.

Thandapani P, O’Connor TR, Bailey TL & Richard S (2013) Defining the RGG/RG Motif. Mol Cell 50, 613–623.

Tran EJ, Zhou Y, Corbett AH & Wente SR (2007) The DEAD-Box Protein Dbp5 Controls mRNA Export by Triggering Specific RNA:Protein Remodeling Events. Mol Cell 28, 850–859.

Vagner S, Rüegsegger U, Gunderson SI, Keller W & Mattaj IW (2000) Position-dependent inhibition of the cleavage step of pre-mRNA 3’-end processing by U1 snRNP. RNA 6, 178–88.

Villarreal OD, Mersaoui SY, Yu Z, Masson JY & Richard S (2020) Genome-wide R-loop analysis defines unique roles for DDX5, XRN2, and PRMT5 in DNA/RNA hybrid resolution. Life Sci Alliance 3.

Viphakone N, Voisinet-Hakil F & Minvielle-Sebastia L (2008) Molecular dissection of mRNA poly(A) tail length control in yeast. Nucleic Acids Res 36, 2418.

Wang K, Wang L, Wang J, Chen S, Shi M & Cheng H (2018) Intronless mRNAs transit through nuclear speckles to gain Zexport competence. Journal of Cell Biology 217, 3912–3929.

Wang L, Tang Y, Cole PA & Marmorstein R (2008) Structure and chemistry of the p300/CBP and Rtt109 histone acetyltransferases: implications for histone acetyltransferase evolution and function. Curr Opin Struct Biol 18, 741–747.

Wang Y, Fan J, Wang J, Zhu Y, Xu L, Tong D & Cheng H (2021) ZFC3H1 prevents RNA trafficking into nuclear speckles through condensation. Nucleic Acids Res 49, 10630–10643.

Weis K & Hondele M (2022) The Role of DEAD-Box ATPases in Gene Expression and the Regulation of RNA-Protein Condensates. Annu Rev Biochem 91, 197–219.

West S, Gromak N & Proudfoot NJ (2004) Human 5’→ 3’ exonuclease Xrn2 promotes transcription termination at co-transcriptional cleavage sites. Nature 432, 522–525.

Wickham H (2016) ggplot2: Elegant Graphics for Data Analysis, Springer-Verlag New York. Available at: https://ggplot2.tidyverse.org.

Wier AD, Mayekar MK, Héroux A, Arndt KM & VanDemark AP (2013) Structural basis for Spt5-mediated recruitment of the Paf1 complex to chromatin. Proc Natl Acad Sci U S A 110, 17290–17295.

Will CL & Lührmann R (2001) RNP Remodeling With DExH/D Boxes. Science (1979) 291, 1916–1917.

Wilson BJ, Bates GJ, Nicol SM, Gregory DJ, Perkins ND & Fuller-Pace F V. (2004) The p68 and p72 DEAD box RNA helicases interact with HDAC1 and repress transcription in a promoter-specific manner. BMC Mol Biol 5.

Wittmann S, Renner M, Watts BR, Adams O, Huseyin M, Baejen C, El Omari K, Kilchert C, Heo D-H, Kecman T, Cramer P, Grimes JM & Vasiljeva L (2017) The conserved protein Seb1 drives transcription termination by binding RNA polymerase II and nascent RNA. Nat Commun 8, 14861.

Yague-Sanz C, Vázquez E, Sánchez M, Antequera F & Hermand D (2017) A conserved role of the RSC chromatin remodeler in the establishment of nucleosome-depleted regions. Curr Genet 63, 187–193.

Yamanaka S, Yamashita A, Harigaya Y, Iwata R, Yamamoto M, Szczesny RJ, Drazkowska K, Pastula A, Andersen JS, Stepien PP, Dziembowski A & Jensen TH (2010) Importance of polyadenylation in the selective elimination of meiotic mRNAs in growing S. pombe cells. EMBO J. 29, 2173–81.

Yamashita A, Shichino Y, Tanaka H, Hiriart E, Touat-Todeschini L, Vavasseur A, Ding D-Q, Hiraoka Y, Verdel A & Yamamoto M (2012) Hexanucleotide motifs mediate recruitment of the RNA elimination machinery to silent meiotic genes. Open Biol. 2, 120014.

Yamashita A, Takayama T, Iwata R & Yamamoto M (2013) A novel factor Iss10 regulates Mmi1-mediated selective elimination of meiotic transcripts. Nucleic Acids Res 41, 9680–7.

Yu Z, Mersaoui SY, Guitton-Sert L, Coulombe Y, Song J, Masson JY & Richard S (2020) DDX5 resolves R-loops at DNA double-strand breaks to promote DNA repair and avoid chromosomal deletions. NAR Cancer 2.

Zenklusen D, Vinciguerra P, Wyss J-C & Stutz F (2002) Stable mRNP Formation and Export Require Cotranscriptional Recruitment of the mRNA Export Factors Yra1p and Sub2p by Hpr1p. Mol. Cell. Biol. 22, 8241–8253.

Zhang X-R, Zhao L, Suo F, Gao Y, Wu Q, Qi X & Du L-L (2022) An improved auxin-inducible degron system for fission yeast. G3 Genes|Genomes|Genetics 12.

Z Zhou Y, Zhu J, Schermann G, Ohle C, Bendrin K, Sugioka-Sugiyama R, Sugiyama T & Fischer T (2015) The fission yeast MTREC complex targets CUTs and unspliced pre-mRNAs to the nuclear exosome. Nat Commun 6, 7050.

