## Supplemental Figures for "DEAD-box ATPase Dbp2 mediates mRNA release after 3’-end formation"

### Supplementary Data

#### Aydin et al., Suppl. Figure 1

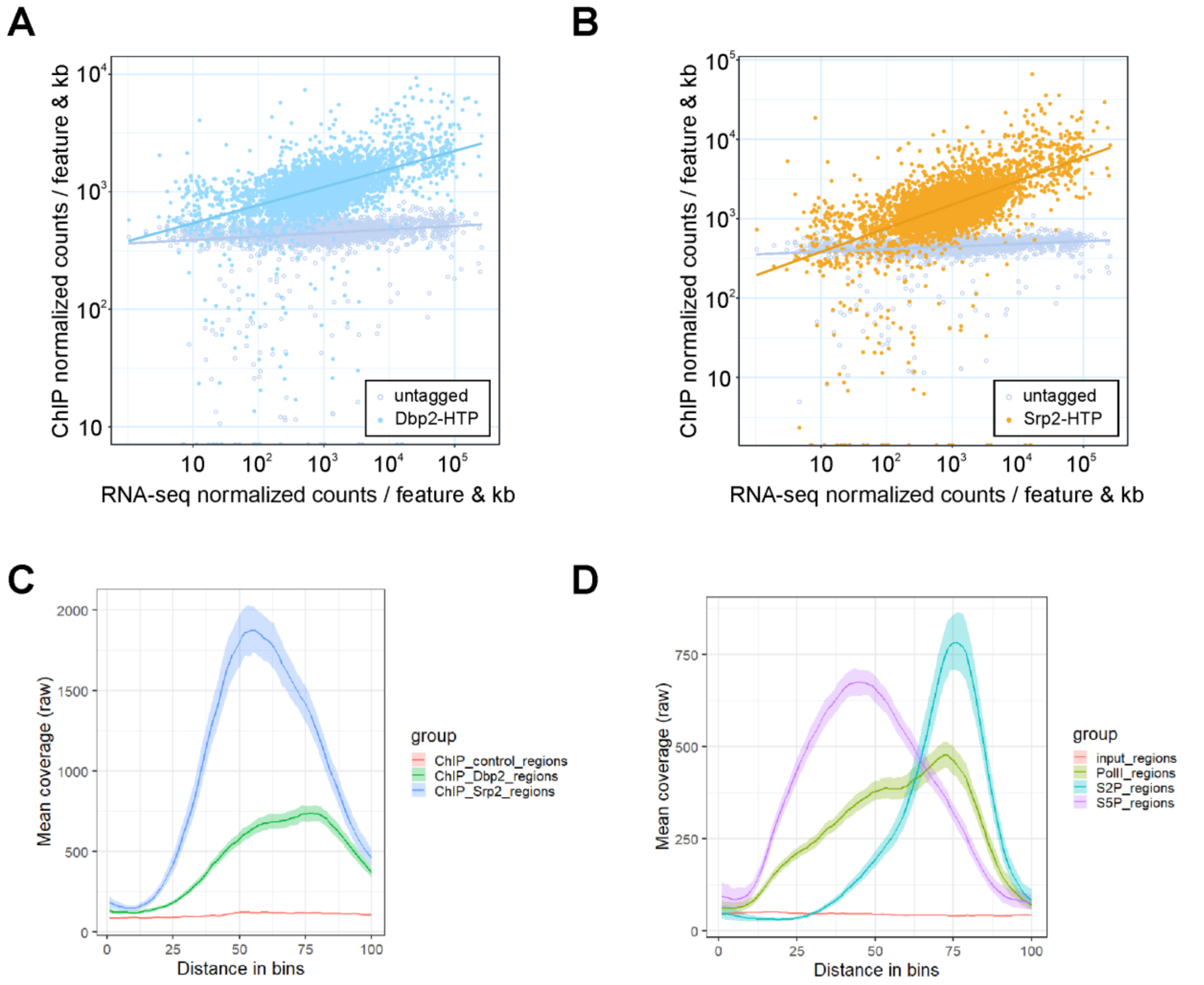


**Supplementary Figure 1.**

**A and B** Integrated counts of Dbp2-HTP (A) and Srp2-HTP (B) ChIP-seq signal across protein-coding genes relative to RNA-seq expression levels in the wild type, given as average counts per feature and kb * 1,000,000 (n = 3). ChIP in the untagged wild type was included as control. Trendlines were fitted using linear regression. RNA-seq expression data is from the untagged isogenic wild type, given as average counts per feature and kb * 1,000,000 (n = 3)**.**

**C and D** Non-scaled curves with confidence intervals for the metagene analysis in Figure 1C, showing metagene analyses of mean ChIP-seq coverage of (**C**) Dbp2-HTP, Srp2-HTP and an untagged wild type control (IgG-ChIP, n = 3) and (**D**) IP against Rpb1, Rpb1-S5P, or Rpb1-S2P, and the corresponding input chromatin (monoclonal antibody ChIP, n = 2) across ribosomal protein genes including 200 bp upstream and 500 bp downstream of the annotated transcription units.

### Aydin et al., Suppl. Figure 2

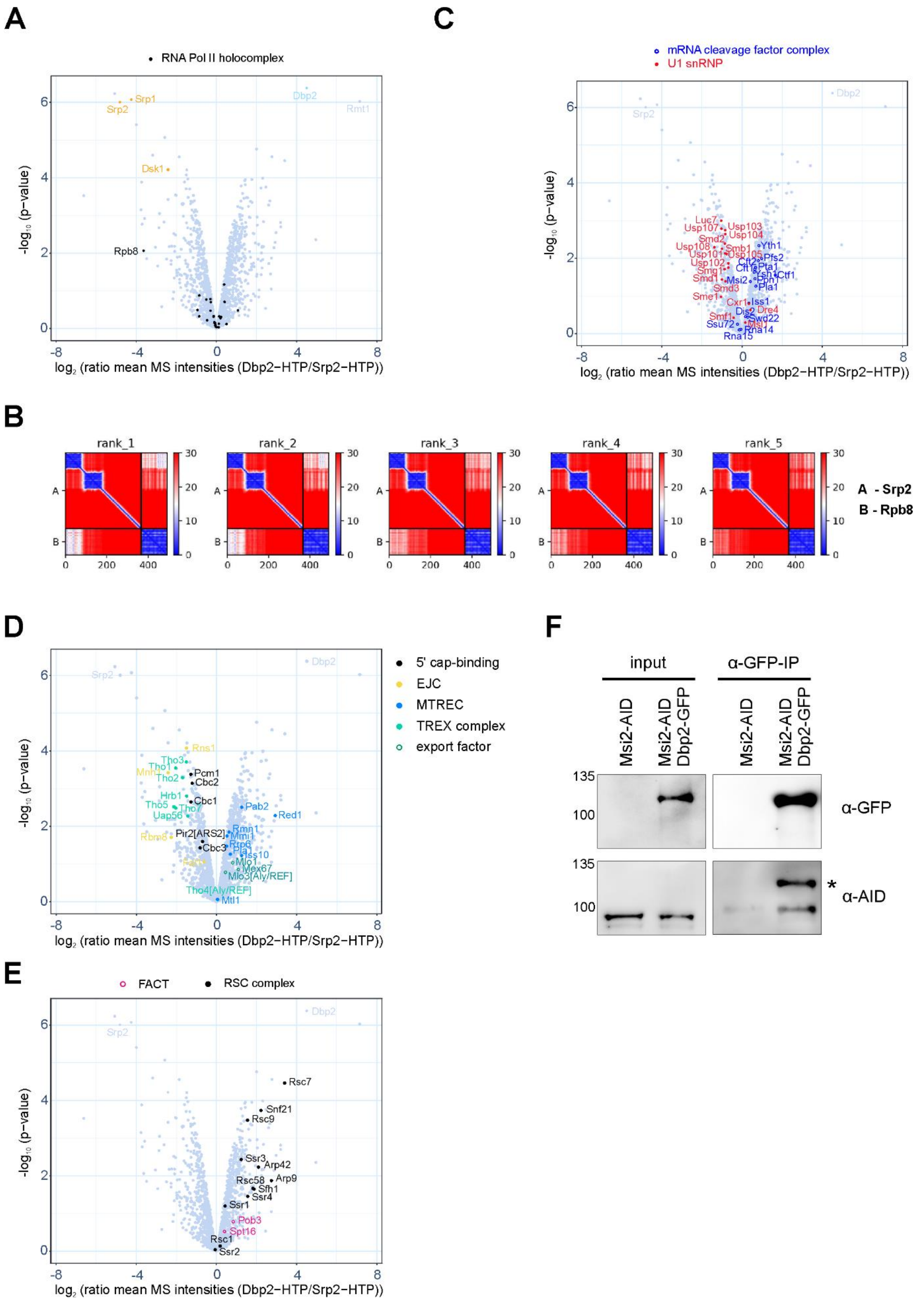

**Supplementary Figure 2**

**A** Mass spectrometry (MS) analysis of the comparative interaction profiling of Dbp2 and Srp2. In the volcano plot, p-values ($-\log_{10}$, moderated Student's t-test) are plotted against the relative enrichment of proteins in the purification of Dbp2-HTP relative to Srp2-HTP based on mean protein intensities ($\log_2$) (n=3). Components of the RNAPII holocomplex (GO:0016591) are marked in black, Srp2 and known interactors in orange, and Dbp2 light blue.

**B** AlphaFold2-multimer prediction of Srp2-Rpb8 interaction, visualised using predicted aligned error (Elfmann & Stülke, 2023). The interaction is predicted by all five default Alphafold2-multimer models, ranked by the confidence with which they predict the interaction. Blue and white shades in the "interaction" quadrants (upper right and lower left) denote low expected position error, indicative of a protein-protein interaction (Evans et al., 2022). Srp2: x-axis amino acid residues 1-365 and y-axis protein A; Rpb8: x-axis amino acid residues 366-490 and y-axis protein B.

**C** MS analysis of the comparative interaction profiling of Dbp2 and Srp2 as in A. Components of the mRNA cleavage factor complex (GO:0005849) and U1 snRNP (GO:0005685) are labelled in blue and red, respectively.

**D** MS analysis of the comparative interaction profiling of Dbp2 and Srp2 as in A. Known nuclear cap-interacting proteins, components of the exon junction complex (EJC), the Mtl1-Red1 core (MTREC), the TREX complex and selected export factors are labelled in black, orange, blue, green, and dark green, respectively.

**E** MS analysis of the comparative interaction profiling of Dbp2 and Srp2 as in A. Components of the RSC complex (GO:0016586) and FACT are labelled in black and magenta, respectively.

**F** Lysates and eluates of a co-immunoprecipitation with GFP-Trap beads from strains expressing Msi2-AID in the presence or absence of GFP-tagged Dbp2 were resolved on SDS-PAGE and analysed by Western blot against GFP and the AID tag. Image is representative of two independent experiments.

### Aydin et al., Suppl. Figure 3

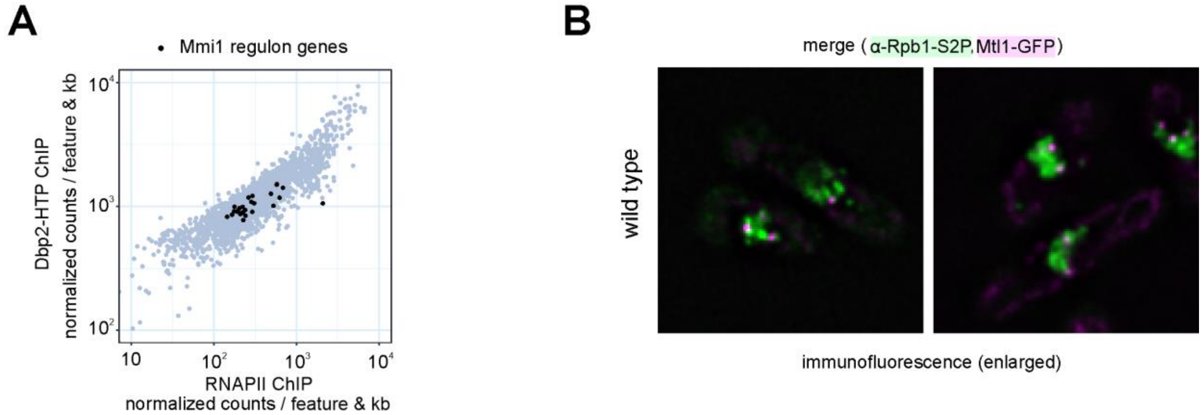


**Supplementary Figure 3**

**A** Integrated counts of Dbp2-HTP ChIP-seq signal across protein-coding genes relative to RNAPII, given as average counts per feature and kb * 1,000,000 (n = 3) as in 1B. Only genes with an RNAPII occupancy above a set threshold of 10 normalized counts per feature and kb per million were included. RNAPII ChIP-seq data from Kecman et al., 2018; GEO: GSE111326 (n = 2). Genes that belong to the "Mmi1 regulon" (Chen et al., 2011) are marked in black.

**B** Enlarged versions of the merged images of the immunofluorescence shown in Figure 3E, with Rpb1-S2P in green and Mtl1-GFP in magenta, without yellow markings.

### Aydin et al., Suppl. Figure 4

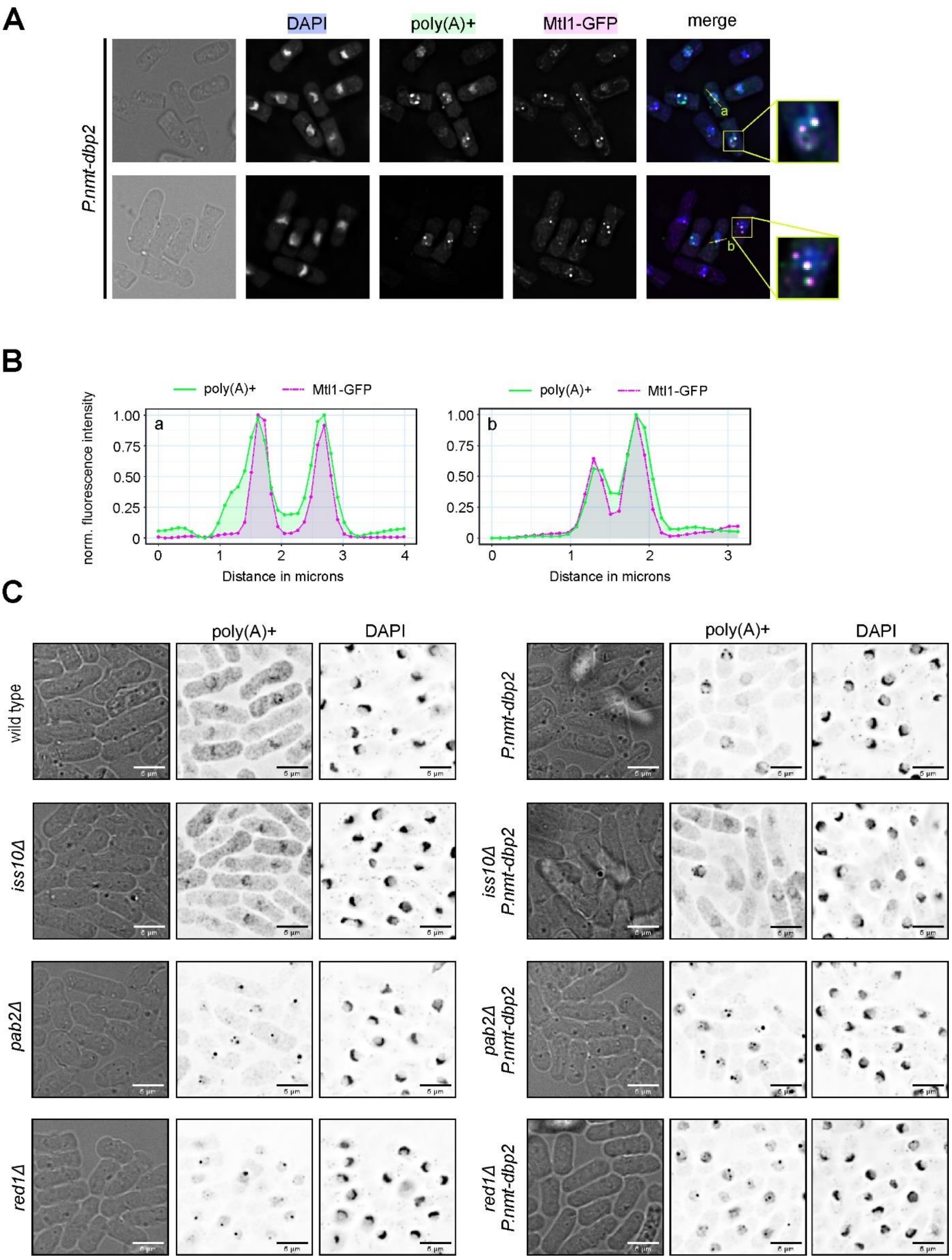


**Supplementary Figure 4**

**A** FISH-IF against poly(A)+ RNA using oligo-d(T)-Cy3 in *P.nmt-dbp2* with the cleavage body marker Mtl1 genomically tagged with GFP. Cells were grown overnight in EMMG, then shifted to YES for 5h to shut off the *P.nmt1* promoter before formaldehyde fixation. The merged channel shows poly(A)+ RNA in green, Mtl1-GFP in magenta and DAPI in blue. Fluorescence intensity profiles were generated along the yellow lines and are shown in D. Images are representative of three independent experiments. Note that in the Mtl1-GFP background, the poly(A)+ RNA signal tends to be more focused than in the untagged strain.

**B** Fluorescence intensity profiles of FISH-IF signal across sites of poly(A)+ RNA accumulation and the adjacent nuclear area as indicated in B. Green line corresponds to the poly(A)+ RNA signal, dashed magenta line to the Mtl1-GFP signal. Fluorescence intensities were normalised to a 0-1 range.

**C** Fluorescence in-situ hybridization (FISH) against poly(A)+ RNA using a Cy3-labelled oligo-d(T) probe and DAPI to stain the DNA within the nucleus. Cells were grown over night in EMMG, then shifted to YES for 5h to shut off the *P.nmt1* promoter before formaldehyde fixation. Fluorescence images were inverted for easier visibility. Images are representative of three independent experiments. A quantitation of the nuclear/cytoplasmic signal distribution is given in 4F.

#### Aydin et al., Suppl. Figure 5

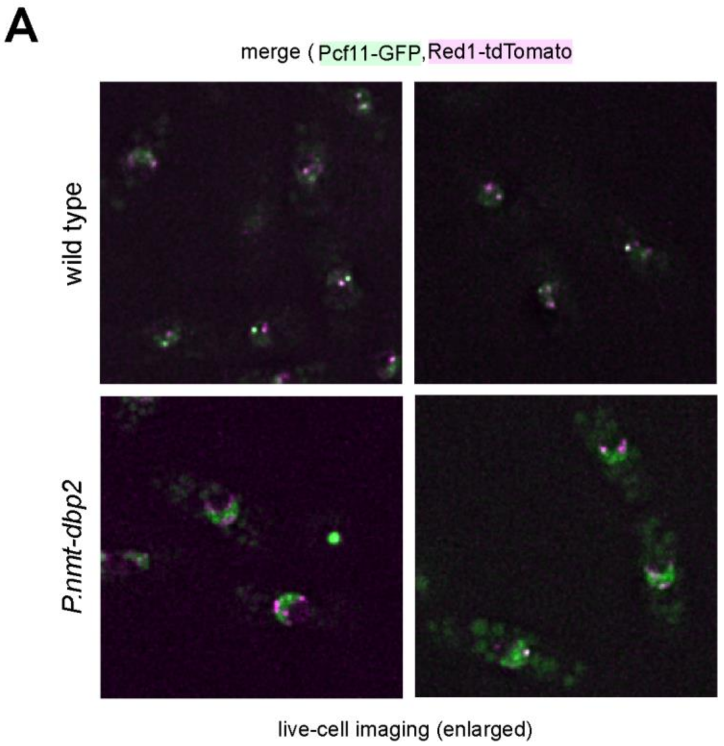


**Supplementary Figure 5**

**A** Enlarged versions of the merged images of the live-cell imaging data shown in Figure 5A, with Pcf11-GFP in green and Red1-tdTomato in magenta, without yellow markings.

### Aydin et al., Suppl. Figure 6

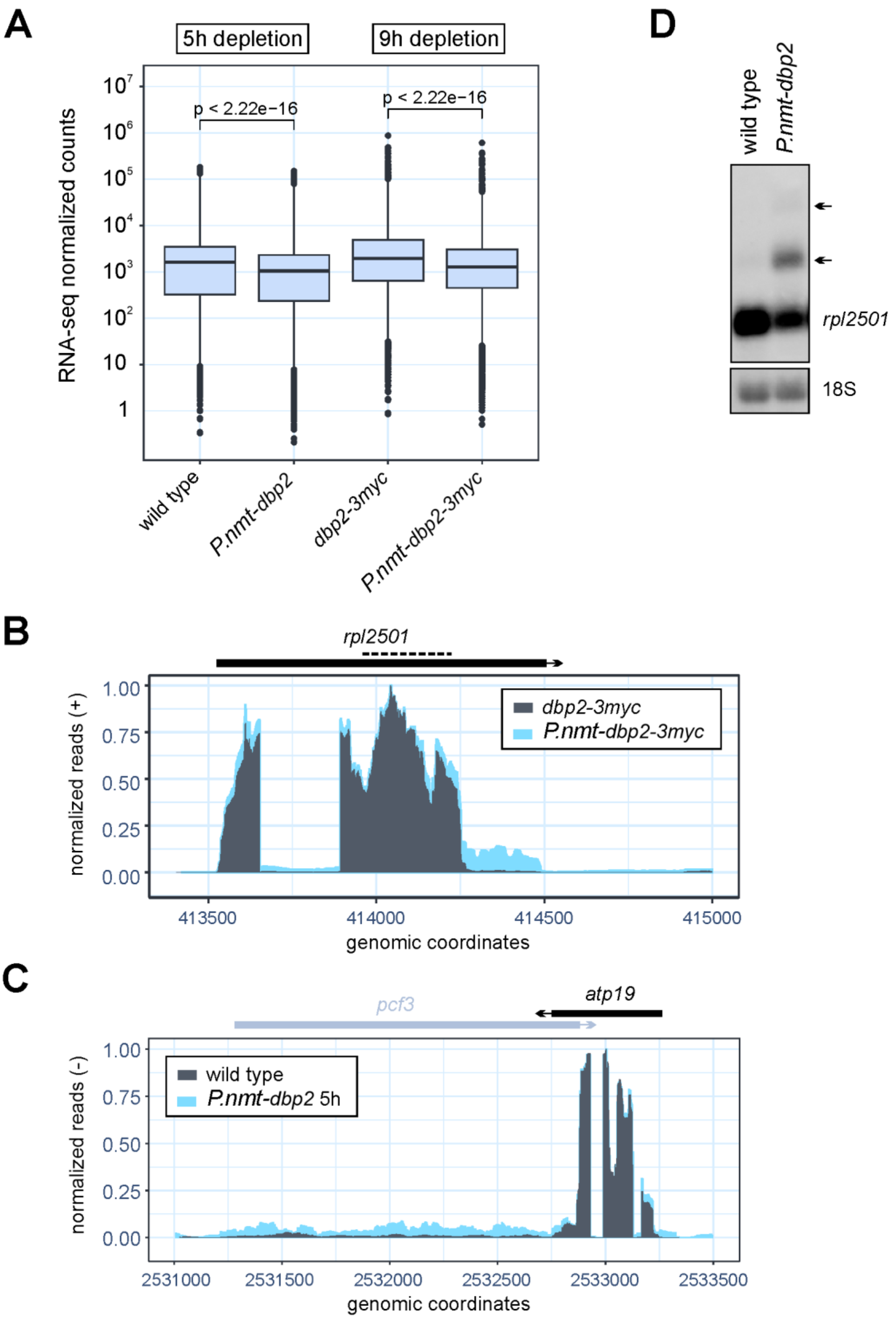


**Supplementary Figure 6**

**A** Mean integrated counts of RNA-seq reads over annotated features in untagged (n = 3) or 3myc-tagged (n = 2) Dbp2 after a 5h or 9h depletion period, respectively. Normalized counts were generated as in Figure 6A. The displayed p-values for the pair-wise comparisons were calculated using the Wilcoxon test.

**B** RNA-seq traces (positive strand) across the region of chromosome II encompassing the *rpl2501* gene with reads for *dbp2-3myc* in grey, for *P.nmt-dbp2-3myc* in light blue. Read counts were normalized to a 0-1 range to visualize relative amounts of 3'-extended transcripts. Cells were grown in EMMG overnight at 30°C, harvested, and cultured in YES medium for 9h prior to RNA isolation and ribodepletion. Dotted line indicates the position of the Northern probe used in D.

**C** RNA-seq traces (negative strand) across the region of chromosome I encompassing the *atp19* gene with reads for the wild type in grey, for *P.nmt-dbp2* in light blue. Read counts were normalized to a 0-1 range to visualize relative amounts of 3'-extended transcripts. Cells were grown in EMMG overnight at 30°C, harvested, and cultured in YES medium for 5h prior to RNA isolation and poly(A) selection.

**D** Northern blot for *rpl2501* mRNA using a strand-specific DIG-labelled RNA probe against the gene body as indicated in B. 18S band stained with methylene blue is shown as loading control. Cells were grown in EMMG overnight at 30°C, harvested, and cultured in YES medium for 5h prior to RNA isolation. Arrows indicate positions of extended transcripts. Images are representative of three independent experiments.

### Aydin et al., Suppl. Figure 7

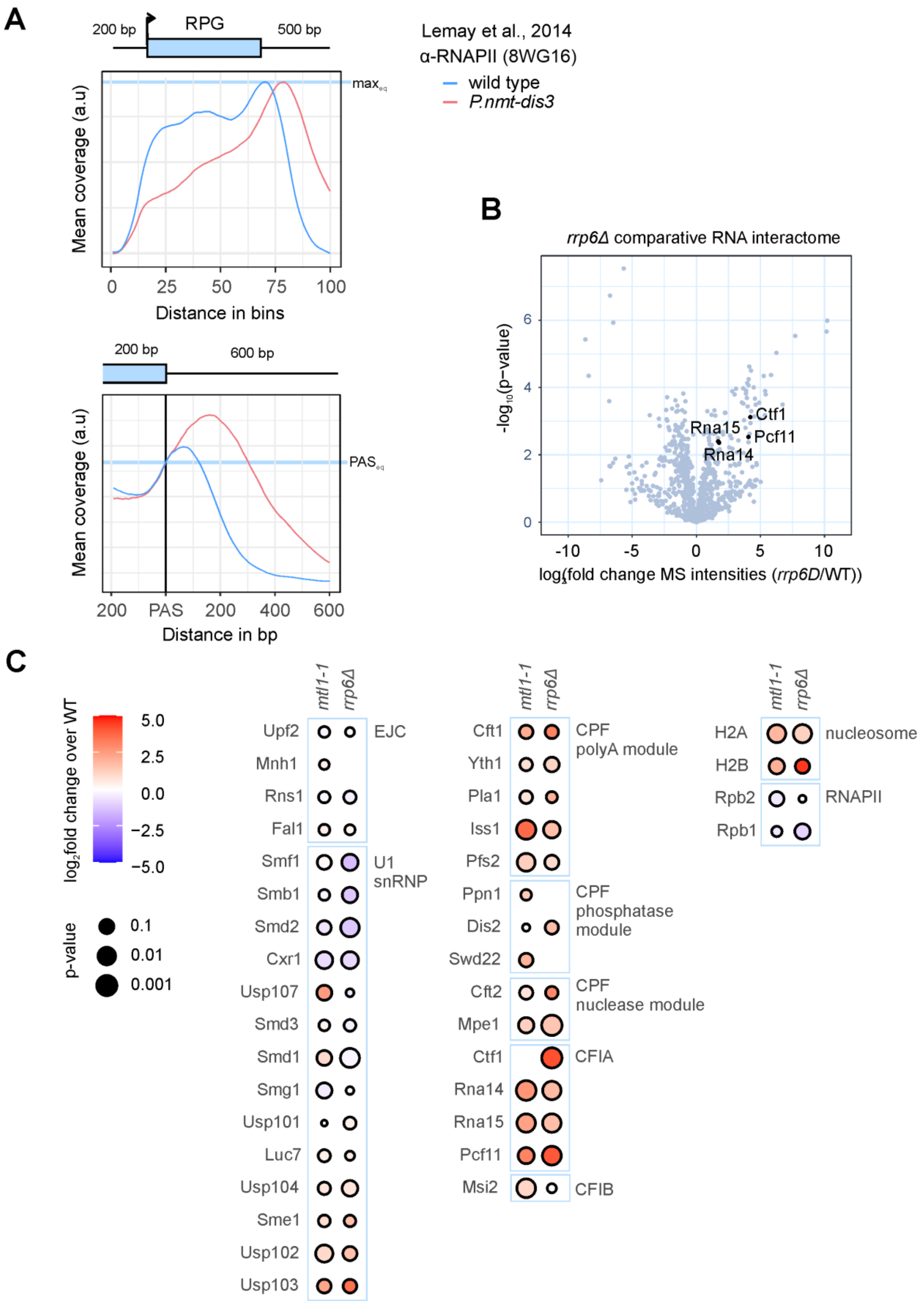

**Supplementary Figure 7**

**A** Metagene analysis of mean RNAPII ChIP-seq signal as in 7C for wild type and *P.nmt-dis3* (n = 2) across ribosomal protein genes (RPGs) including 200 bp upstream and 500 bp downstream of the annotated transcription units (upper panel) or surrounding the polyadenylation and cleavage site (PAS) (lower panel). Mean coverage was adjusted by a constant scaling factor to normalize to maximal peak height (upper panel) or RNAPII levels at the PAS (lower panel) for an easier comparison of curve shapes. Schematic of the gene above the left panel corresponds to an RPG of median length. Within the metagene, the positions of transcription start and end sites are distributed around the given coordinate because of the varying feature compression depending on gene length. Data from (Lemay et al., 2014), ArrayExpress accession E-MTAB-2237; cells were grown in EMMG and Dis3 depleted by addition of thiamine for > 12h.

**B** MS analysis of a comparative poly(A)+ RNA interactome capture experiment for the nuclear exosome mutant *rrp6Δ*. In the volcano plot, p-values (−log, moderated Student's t-test) are plotted against the fold change of mean MS intensities ($\log_2$) of proteins recovered from the oligo(dT) pull-downs of UV-crosslinked samples (3 $J/cm_2$) for wild type and *rrp6Δ* (n = 3). Data from (Kilchert et al., 2020), ProteomeXchange accession PXD016741. CF1A components are enriched on poly(A)+ RNA in the exosome mutant.

**C** Enrichment of CPAC components, various splicing factors, and representative RNAPII and nucleosome components on poly(A)+ RNA in the nuclear exosome mutants *mtl1-1* and *rrp6Δ* based on the MS analysis of comparative poly(A)+ RNA interactome capture experiments as in B. Colour indicates the $\log_2$ fold change (mutant over wild type), circle size the p-value (moderated Student's t-test). Data from (Kilchert et al., 2020), ProteomeXchange accession PXD016741; cells were grown in EMMG and labelled with 4-thiouracil for 4.5h prior to crosslinking.
