## Supplemental File: FISH quant ImageJ script for "DEAD-box ATPase Dbp2 mediates mRNA release after 3’-end formation"

// quantification of RNA export (oligo-dT-FISH) using transmitted light and DAPI as reference channels
// stick to naming conventions as indicated to keep compatibility with downstream R scripit

// select cellular content based on thresholding
	// --> select image stack
	// choose transmitted light image at low range so that white rim shows up (for me, usually 2-3)
		  Dialog.create("Set variable");
		  	Dialog.addNumber("DIC channel:", 1);
		  	Dialog.addNumber("z slice:", 3);
		  	Dialog.show();
		  	channel = Dialog.getNumber();
		  	zSlice = Dialog.getNumber();
		  	print(channel);
		  	print(zSlice);
		run("Duplicate...", "duplicate channels=channel slices=zSlice");
		run("Bandpass Filter...", "filter_large=40 filter_small=3 suppress=None tolerance=5 autoscale saturate");
		run("Median...", "radius=2");
		run("Threshold...");
	// threshold manually so that cell content is selected and clearly separated from surroundings
		
//// BREAKPOINT ////	
		run("Invert");
		
					// optional: if necessary, fill segmentation gaps along the cell walls manually, to do so, invert im age, use manual selection tool and clear spaces
					// eroding or dilating one round can help; reinvert before continuing 
						run("Convert to Mask");
						run("Options...", "iterations=1 count=1 do=Erode");
						run("Options...", "iterations=1 count=1 do=Dilate");
						
// on final image, select cell bodies via analyze particles (area around thirty, circulartiy around 0.5); if necessary, use results table to detrermine best settings
	run("Duplicate...", " ");
	run("Analyze Particles...", "size=7.00-60.00 circularity=0.10-1.00 show=Masks display exclude clear add in_situ");

//// BREAKPOINT /////
// renaming ROIs to cell1 - cellXY
// before running: check selected ROIs and delete any obvious junk
		roiManager("UseNames", "false");
		for (i = 0; i < roiManager("count") ; i++) {
		    roiManager("Select", i);
		    roiManager("Rename", "cell"+i+1);
		    Roi.setGroup(4);
		}

//// BREAKPOINT /////
// selects nuclei on DAPI channel; here we use a z projection across the entire stack
	// --> select image stack ///
			Dialog.create("Set variable");
		  	Dialog.addNumber("DAPI channel:", 3);
		  	Dialog.show();
		  	channel = Dialog.getNumber();
			run("Duplicate...", "duplicate channels=channel");
			run("Z Project...", "projection=[Average Intensity]");
			run("Duplicate...", " ");
			run("Threshold...");
		// set threshold manually to select DAPI-stained nuclei
			
//// BREAKPOINT ///			
			run("Convert to Mask");
			run("Duplicate...", " ");
			// because DAPI does not fill the entire nucleus (compare heat shock mRNA retention signal), we dilate the nuclear signal
			// if stuff touches, select particles first, clear ROIs on empty image, dilate, then re-select
			run("Options...", "iterations=1 count=1 do=Dilate");
			// on final image, select nuclei via analyze particles (circularity around 0.8); use results table to adjust, if necessary
		run("Analyze Particles...", "size=1-9.00 circularity=0.50-1.00 show=Masks display exclude add in_situ");

//// BREAKPOINT /////
// Overlapping ROIs select and pairwise renaming
/// this will work on a set of ROIs where cells are labelled cell1 to cellXY (in ascending order without gaps!) and ROIs for nuclei have already been added
/// nuclei ROIs can be in random order and with random names; nuclei without cells and cells without nuclei are fine --> will be removed  
/// you may want to save your ROIs before you start the clean-up


	// Overlapping ROIs selection: finds nuclei that overlap with each cell and renames them accordingly
	//for cells with two nuclei, both will get the same name (these will be joined later based on the name duplication)
			cellArray = newArray();  
							for (i = 0; i < roiManager("Count"); i++){
								roiManager("Select", i);
								if (Roi.getGroup()==4) {
									cellArray = Array.concat(cellArray, i);
								} 
							}
							cellNumber = cellArray.length;
							print("there are ", cellNumber, " cells");
	for (i = 0; i < cellNumber; i++){ 
		for (j = cellNumber; j < roiManager("Count"); j++){
		   if (i != j){
		   	   	roiManager("Select", newArray(i,j));
		   		roiManager("AND");
		    
				if (selectionType()>-1) {
					print (i,j," do intersect");
					roiManager("Deselect");
					roiManager("Select", j);
					roiManager("Rename", "nuc"+i+1);
					Roi.setGroup(1); /// all nuclei that were assigned to cells
					roiManager("Deselect");
					roiManager("Select", i);
					Roi.setGroup(2); /// all cells that were assigned to nuclei
		   		} 
		   }
		}
	}
					/// all ROIs that do not have a matching overlapping ROI were not assigned to a group and are discarded 				
						indexList = newArray();
						for (i = 0; i < roiManager("Count"); i++){
							roiManager("Select", i);
							if (Roi.getGroup()==0) {
								indexList = Array.concat(indexList, i);
								print(i, "had no matching ROI");
								;
							} else {
							if (Roi.getGroup()==4) {
								indexList = Array.concat(indexList, i);
								print(i, "had no matching ROI");
							} 
							}
						}
						roiManager("Select", indexList);
						roiManager("Delete");

	/// merges nuclei mapping to the same nucleus, which have identical names
	/// retrieving and comparing names is time-consuming --> this sorts first and only compares with neighbour

			/// counts number of cells (in ROI group 2) and stores as variable
			cellArray = newArray();  
							for (i = 0; i < roiManager("Count"); i++){
								roiManager("Select", i);
								if (Roi.getGroup()==2) {
									cellArray = Array.concat(cellArray, i);
								} 
							}
							cellNumber = cellArray.length;
							print("there are ", cellNumber, " cells");
				/// loops through nuclei and merges those with the same name; adds merged ROI to manager and deletes original ROIs			
				roiManager("Sort");
				indexList = newArray();
				for (i = cellNumber; i < roiManager("Count")-1; i++){ 
					  		roiManager("Select", i);
					   		roiName = Roi.getName();
					   		roiManager("Deselect");
					   		roiManager("Select", i+1);
					   		roiName2 = Roi.getName();
					   	   	if (roiName == roiName2) {
					   	   		print (i,i+1," are assigned to the same cell as", roiName);
					   	   		roiManager("Deselect");
					   	   		roiManager("Select", newArray(i,i+1));
					   			roiManager("OR");
					   			roiManager("Add");
					   			roiManager("Select", roiManager("Count")-1); 
					   			roiManager("Rename", roiName);
					   			indexList = Array.concat(indexList, i);
					   			indexList = Array.concat(indexList, i+1);
					   	   	}
					   }
					   	roiManager("Select", indexList);
						roiManager("Delete");

//// optional BREAKPOINT /////
/// do a visual check whether ROIs were cleaned up properly
// afterwards, uses XOR command on nuc1 & cell1 to generate ROI for cytoplasm, rename cyto1-XY
	// sort alphabetically
	roiManager("Sort");
		/// counts number of cells and nuclei and writes to log
		/// you may want to run the count once without the follow-up to make sure the numbers match
		/// BUT: you will need to re-run along with the XOR and relabel blocks  to generate the cellNumber variable
		cellArray = newArray();  
		nucArray = newArray();
					for (i = 0; i < roiManager("Count"); i++){
						roiManager("Select", i);
						if (Roi.getGroup()==2) {
							cellArray = Array.concat(cellArray, i);
						} 
					}
					cellNumber = cellArray.length;
					print("there are ", cellNumber, " cells and ", roiManager("Count")-cellNumber, " nuclei");

					/// runs XOR on sorted ROIs to generate cytoplasm ROI 
					for (i = 0; i < cellNumber; i++) {
					 	roiManager("Select", i);
					  	roiManager("Select", newArray(i,i+cellNumber));
						roiManager("XOR");
						roiManager("Add");
					}
								/// relabel all ROIs 
								// cells
									for (i = 0; i < cellNumber ; i++) {
									    roiManager("Select", i);
									    roiManager("Rename", "cell"+i+1);
									}	
									// to label nucleus
									for (i = cellNumber; i < cellNumber*2; i++) {
									    roiManager("Select", i);
									    roiManager("Rename", "nuc"+i+1-cellNumber);
									}	
									// to label cytoplasm
									for (i = cellNumber*2; i < cellNumber*3; i++) {
									    roiManager("Select", i);
									    roiManager("Rename", "cyto"+i+1-cellNumber*2);
									}	

///	optional block --> alternatively, select with ctrl-A and save manually
								// select all ROIs, export  via more... / save RoiSet
										// selects all ROIs
										count = roiManager("count");
										array = newArray(count);
										  for (i=0; i<array.length; i++) {
										      array[i] = i;
										  }
										roiManager("select", array);
										// roiManager("Save", "folder/filename");
//// BREAKPOINT /////									
// quantify FISH signal using ROIs
// select image stack
			Dialog.create("Set variable");
		  	Dialog.addNumber("FISH channel:", 2);
		  	Dialog.show();
		  	channel = Dialog.getNumber();
			run("Duplicate...", "duplicate channels=channel");
		//run("Brightness/Contrast...");
		run("Z Project...", "projection=[Average Intensity]");
		run("Set Measurements...", "area mean min shape integrated display redirect=None decimal=5");
		run("Clear Results");
		// select all ROIs, hit measure on AVG image
		roiManager("Show All");
		roiManager("Measure");
		// save results
		// for documentation, make a screen shot of ROIs on both AVG-DAPI and AVG-FISH
